# Lydicamycins Induce Morphological Differentiation in Actinobacterial Interactions

**DOI:** 10.1101/2024.06.28.600750

**Authors:** Scott A. Jarmusch, Morten D. Schostag, Zhijie Yang, Jinglin Wang, Aaron J.C. Anderson, Tilmann Weber, Ling Ding

## Abstract

*Streptomyces* are major players in soil microbiomes, however, interactions involving *Streptomyces* and other actinobacteria are rarely described. The complex developmental cycle of actinobacteria necessitates a multi-omics approach to unravel the web of information. This study resulted from the observation of induced morphogenesis between two environmental isolates from the same site, *Kitasatospora* sp. P9-2B1 and *Streptomyces* sp. P9-2B2. When co-cultivated on potato dextrose agar (PDA), P9-2B2 induced a wave-like sporulation in strain P9-2B1. Using mass *s*pectrometry imaging, we revealed that a suite of lydicamycins were present in this induced sporulation zone. Lydicamycin deficient mutants were generated using CRISPR Base-Editing and the inducible sporulation ceased, confirming their role in triggering morphological differentiation. In agar diffusion assays, pure lydicamycin was inhibitory when added concurrently with bacterial inoculation and induced sporulation with delayed addition. Subsequent testing of additional environmental isolates resulted in the same inducible sporulation wave phenomenon, including *Streptomyces coelicolor* M145 and M1146. Using transcriptomics, we observed the differential expression of genes related to early aerial mycelium development four days into cocultivation and the transitional genes responsible for development of spores on day 9. Along with these differentially expressed genes, we also observed numerous overall stress responses, specifically cell envelope stress responses. These findings uncovered actinobacteria interactions mediated by lydicamycins, pointing to a potential role of certain groups of bioactive metabolites in nature.

**Importance:** Shifting away from an antibiotic discovery mindset, uncovering the chemical ecology of secondary metabolites is key to maximizing their biotechnological application. The reduced complexity of dual cultures allows for in depth analysis and investigation of these interactions via multi-omics provides orthogonal data leading to more robust conclusions. This study provides insight into the role of lydicamycins in dual cultures with other actinobacteria and provides an integral roadmap for future chemical ecology work between microorganisms, especially through the use of mass spectrometry imaging.

## Introduction

*Streptomyces* are one of the most widely utilized natural sources, principally for their prolific production of ∼70% of clinical antibiotics (1, 2). The attention to this veritable ‘arms race’ has diverted resources towards drug discovery, whereas in contrast, the wider ecological influence of *Streptomyces* secondary metabolites (SMs) has severely lagged behind, despite of few reported signaling molecules, *e.g.* γ-butyrolactones(3), governing the production of other SMs or transition of life stage. *Streptomyces* serve vital roles in soil microbiomes, evidenced by their diverse ecological distribution and their SMs are the ‘language’ they communicate with friends and foes (4, 5). While interesting observations have been found for microbial inter- and intra-kingdom interactions, such as *Streptomyces*-fungus(6) and *Streptomyces*-*Bacillus subtilis* interactions(7), respectively, a large gap exists in understanding the role SMs play in actinobacteria interactions.

Sporulation is a common phenotype change observed in *Streptomyces* interactions(8), yet tracing the responsible SMs has rarely been studied. Goadsporin was the first SM described to cause sporulation in *Streptomyces* (9). A major challenge is linking the phenotype change to the secondary metabolite; identifying the metabolite from the earliest stages of analysis is the bottleneck allowing for efficient linkage of SMs and ecology. Mass Spectrometry Imaging (MSI) offers attractive solutions to resolving this spatial problem(10, 11) through direct visualization of metabolites on cocultivations (12). In *Streptomyces*-actinobacteria studies, Traxler et al. set the primer for evaluating actinobacterial cocultivations, leading to the discovery of new acylated desferrioxamines (13), and many studies have followed thereafter proving MSI on cocultures is a faster route to discovery of novel compounds with cocultures.

During a screening for *Streptomyces*-actinobacteria antibiosis, one environmental strain (*Streptomyces* sp. P9-2B2) induced a wave of sporulation in a receiver strain (*Kitasatospora* sp. P9-2B1). Using a combination of MSI, molecular genetics and biological testing, we determined that P9-2B2 produced a suite of antibiotic lydicamycins responsible for the induced morphogenesis. The temporal production of lydicamycins was evaluated using LC-MS/MS and feature based molecular networking. Finally, we performed transcriptomics analysis on a coculture between *Streptomyces coelicolor* M1146 and P9-2B2 to determine the differentially expressed genes during lydicamycin exposure.

## Materials and methods

### Selective isolation and genomics

All environmental isolates used in this study were isolated from soil collected in 2020 from the UNESCO World Heritage Site, Jægorsborg Deer Park (Dyrehaven), Denmark. Selective isolation using standard techniques were carried out to isolate actinobacteria and specifically Streptomycetes (14, 15). The genome sequence data were analyzed with the Type Strain Genome Server (TYGS), a free bioinformatics platform available under https://tygs.dsmz.de, for a whole genome-based taxonomic analysis. Both P9-2B1 and P9-2B3 were identified as isolates of *Kitasatospora papulosa* (dDDH of 94.2% and 93.4% compared to *Kitasatospora papulosa* NRRL B-16504) while P9-2B2 and P9-2B4 appeared as *Streptomyces platensis* (dDDH of 74.1% and 76.2% compared to *Streptomyces platensis* DSM40041). More detailed methods and phylogenetic trees can be found in supplementary data (Figs. S1-4). The genome sequences of *Kitasatospora* sp. P9-2B1 and *Streptomyces* sp. P9-2B2 were deposited into BioProject PRJNA985726. *Streptomyces coelicolor* M145 and M1146 were obtained from Prof. Mervyn Bibb, John Innes Centre, Norwich.

### Actinobacteria cocultivation and monocultures

All spore stocks used in this study were standardized to 10^5^ CFU/mL, 10 µL of inoculum was added for each isolate, and all cocultivations were carried out on BD Difco Potato Dextrose Agar (PDA). All cocultivations were spaced 1.5 cm apart. Inocula used for MSI were incubated at 30 °C for 7 days and then prepared for imaging. Inocula used for timelapse imaging were placed in a 30 °C incubator room with a Reshape Timelapse Imager (Reshape, Copenhagen, DK) and one image was taken every 60 minutes for 7-10 days depending on experiment length. Experiments involving pure lydicamycin (Santa Cruz Biotechnology Inc.): bacterial spore stocks were plated and pre-incubated for 3 days and then pure metabolites were added to an agar well 1.5 cm away. Pre-incubation was carried out to (1) mimic the time point lydicamycin production is seen in P9-2B2 monocultures and (2) to prevent inhibitory effects of the metabolite. P9-2B2 monocultures for temporal SM production: 10 PDA plates with three 10 µL spots were incubated, where every day one plate was removed, one colony was imaged using a Leica S9i digital microscope and extracted as described below. The exact same procedure was carried out with ISP2 plates but only sampling on days 1, 2, 3, 4, 7 and 10. Supplementary microscopic images were acquired by a Zeiss SteREO Discovery.V12 stereo microscope, equipped with a Zeiss Axiocam 702 mono digital camera (Carls Zeiss AG, Oberkochen, Germany).

### Inactivation of the lydicamycin BGC

In order to inactivate the core biosynthetic gene *lyd60* (NCBI Genbank accession: WP_290354452.1), coding for the first modules of the lydicamycin PKS, a STOP codon via CRISPR-BEST base editing (16) (*lyd*60^STOP^) was designed using the online tool CRISPy-web (17). pCRISPR-cBEST plasmid was linearized by NcoI and NEBuilder HiFi DNA Assembly Master Mix was used to insert the *lyd*60^STOP^ deactivation, resulting in pCRISPR-cBEST/*lyd60*. Subsequently, Mach1 T1 Phage-Resistant Chemically Competent *E. coli* was transformed with the plasmid. The plasmid was re-isolated and transferred into chemically competent *E. coli* ET12567/pUZ8002. The *E. coli*-*Streptomyces* conjugation experiment was conducted according to the standard protocol (1) and *Streptomyces* mutants were confirmed by Sanger sequencing.

### Sample preparation and Mass Spectrometry Imaging

10 µL glycerol spore stocks of each microbe were plated onto 4 mm thick PDA agar for all MSI experiments. Sample preparation was adapted from Yang et al (18). Upon observation of induced sporulation in the receiver strain, colonies were excised using a scalpel and placed on a Bruker IntelliSlide coated with a layer of glue applied with a glue pen. Samples were subsequently dried for 2-4 hours at 35°C followed by matrix application (40 mg/mL DHB, 50:50:0.1% H_2_O:MeOH:TFA) for 15 passes using HTX-Sprayer (HTX Imaging, Chapel Hill, NC, USA). Samples were further dried at 35°C for 1 hour prior to MSI. The samples were then subjected to timsTOF flex (Bruker Daltonik GmbH, Bremen, GE) mass spectrometer for MALDI MSI acquisition in positive MS scan mode with 20 µm raster width and a mass range of 100-2000 *m/z*. Calibration was done using red phosphorus. The settings in the timsControl were as follow: Laser: imaging 20 µm, Power Boost 3.0%, scan range 16 µm in the XY interval, and laser power 70%; Tune: Funnel 1 RF 300 Vpp, Funnel 2 RF 300 Vpp, Multipole RF 300 Vpp, isCID 0 eV, Deflection Delta 70 V, MALDI plate offset 100 V, quadrupole ion energy 5 eV, quadrupole loss mass 100 m/z, collision energy 10 eV, focus pre TOF transfer time 75 µs, pre-pulse storage 8 µs. All data was analyzed using Bruker SciLS (2021b Pro) and data was normalized to the root mean squared.

### MS-based Metabolomics

All extracts for LC-MS were generated using an agar plug method (19). Liquid chromatography was performed on an Agilent Infinity 1290 UHPLC system. 1 µL extract was injected onto an Agilent Poroshell 120 phenyl-C6 column (2.1 × 150 mm, 1.9 μm) at 40°C using CH_3_CN and H_2_O, both containing 20 mM formic acid. Initially, a linear gradient of 10% CH_3_CN/H_2_O to 100% CH_3_CN over 10 min was employed, followed by isocratic elution of 100% CH_3_CN for 2 min. Then, the gradient was returned to 10% CH_3_CN/H_2_O in 0.1 min and finally an isocratic condition of 10% CH_3_CN/ H_2_O for 1.9 min, all at a flow rate of 0.35 min mL^-1^. HRMS data was recorded in positive ionization on an Agilent 6545 QTOF MS equipped with an Agilent Dual Jet Stream electrospray ion (ESI) source with a drying gas temperature of 250°C, drying gas flow of 8 min L ^-1^, sheath gas temperature of 300°C and sheath gas flow of 12 min L ^-1^. Capillary voltage was 4000 V and nozzle voltage was set to 500 V. The HRMS data were processed and analyzed using Agilent MassHunter Qualitative Analysis B.07.00. HPLC grade solvents (VWR Chemicals) were used for extractions while LCMS grade solvents (VWR Chemicals) were used for LCMS.

Raw data was converted to .mzML using MSConvert (ProteoWizard) and preprocessed using MZmine 3 (20). Molecular networking was all completed within the GNPS platform (21), which includes: Feature Based Molecular Networking (22). Visualization of the molecular networks was completed using Cytoscape 3.8.2. (23).

### RNA extraction and transcriptomic analysis

*Streptomyces* sp. P9-2B2 (producer) was inoculated in the center of the agar plate with *Streptomyces coelicolor* M1146 replicates (*n* = 4) surrounding it at a distance of 1.5 cm. The bacteria were grown at 30 °C and at each time point (day 2, 4, 7 and 9), half of the colony facing the center was collected using a 10 µl inoculation loop and transferred to 2 ml cryovial containing 100 µl phosphate-buffer and 200 µl RNAProtect (QIAGEN N.V., Venlo, The Netherlands). Samples were then centrifuged at 8000g for 5 min, the supernatant was discarded and samples were snap frozen in liquid nitrogen and stored in the −80 °C freezer, prior to RNA extraction.

RNA extraction was performed using the RNeasy Mini kit following the manufacturer’s instructions, with the addition of beat beading with the FastPrep (MP Biomedicals, Santa Anna, USA) for 30 sec at 4500 RPM with 0.1 mm glass beads. DNA was removed with TurboTM DNase kit (Thermo Fisher Scientific). RNA concentration was estimated with Qubit Thermo Fisher Scientific using the High Sensitivity RNA kit (concentrations ranging from 62-781 ng µL-1), and RNA integrity was assessed on an Agilent Bioanalyzer 2100 using Agilent RNA 6000 nano kit. All samples had an RNA integrity number (RIN), with an average of 5,5 (range 2,8-9,0). Ribosomal RNA depletion, library preparation, and sequencing were performed at Novogene Co., Ltd on a NovaSeq PE150 platform. A detailed description of RNA-seq analysis can be found in supplementary methods.

## Results

### Diffusible metabolites induce sporulation of *Kitasatospora* sp. P9-2B1

While investigating actinobacteria interactions, pairwise co-cultivation was carried out and it revealed that *Streptomyces* sp. P9-2B2 caused sporulation of *Kitasatospora* sp. P9-2B1. After 7 days of growth, it was observed that P9-2B1 had a zone of sporulation emanating from P9-2B2, indicating the potential of a diffused metabolite causing morphogenesis. The observed morphogenesis of P9-2B1 occurred in three distinct phases starting from the edge closest to P9-2B2 (Fig. 1A): (1) substrate hyphae first changed in pigmentation, from pale to bright yellow; (2) production of aerial hyphae (white); (3) production of spores (greenish gray). Although the mechanism behind sporulation and the transcriptional effects of sporulation are well understood, metabolites which cause sporulation are essentially unknown, thus peaking our excitement.

**Fig. 1.**
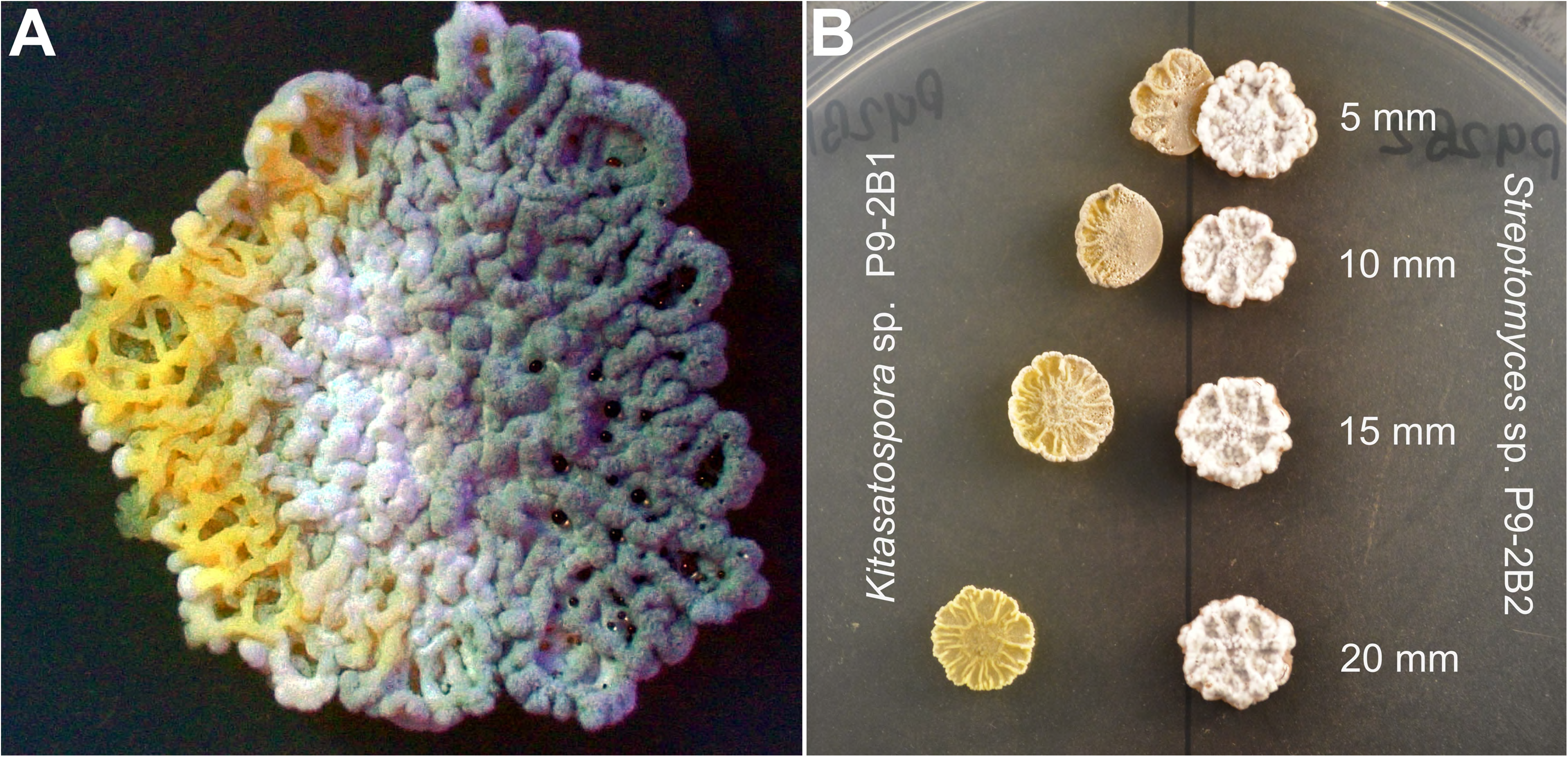
(A) Observed differentiation of *Kitasatospora* sp. P9-2B1 on PDA after 8 days in cocultivation with *Streptomyces* sp. P9-2B2 (cultured on the right side of P9-2B1). (B) Distance based assay showing the diffusible zone of sporulation induced by secondary metabolite production by *Streptomyces* sp. P9-2B2.

Due to the ‘wave-like’ nature of the morphogenesis, we proposed an agar diffusible metabolite produced by P9-2B2 was causing the phenotype change in P9-2B1. To initially confirm the hypothesis, a distance-based assay was carried out where P9-2B1 was incrementally (0.5 cm) moved away from the producer, P9-2B2. As P9-2B1 is distanced farther away, the inducible morphogenesis disappears (Fig. 1B). This indicates that diffusible metabolites are indeed responsible for the phenotype change, similar to previously reported observations of actinobacteria induced sporulation (8). Furthermore, when the P9-2B1 macro-colonies are in very close proximity (5-10 mm), their growth is visibly affected at the nearest point to P9-2B2. In contrast, the colony at 15 mm distance is more comparable to those at 20-25 mm in overall shape, yet aerial hyphae production has begun.

### Mass Spectrometry Imaging illuminates’ sporulation caused by lydicamycin production

Traditionally, researchers have attempted to characterize chemistry derived from cocultures using standard extraction techniques (12, 19). Therefore, our first attempt to identify the responsible metabolites was via agar plug extracts taken from three distinct locations and analyzed using LC-MS: (1) P9-2B2, P9-2B1 (non-sporulating area) and P9-2B1 (sporulated area). No candidate metabolites were differentially observed when comparing the base peak chromatograms nor did the chemistry of P9-2B1 change in these sporulated areas (Fig. S5). The lack of identifying any viable SMs with agar plugs and LCMS lead us to use Mass Spectrometry Imaging (MSI) to directly visualize the spatial metabolome.

Using an adapted method based on previous studies focusing on imaging microbial interactions on agar (18, 24, 25), actinobacteria interactions were carried out on PDA, excised and prepared for MSI. Due to the long growth time required to observe morphogenesis (6-9 days) and substrate mycelium ‘digging’ into the agar causing severe agar splitting and subsequent flaking, we used 2-4 mm agar instead of the standard 1-2 mm, still allowing for proper sample adherence to glass slides as well as minimal air bubbles and flaking. The half-sporulated sample of P9-2B1 (Fig. 2A) as well as two monocultures were analyzed using MALDI-MSI.

**Fig. 2.**
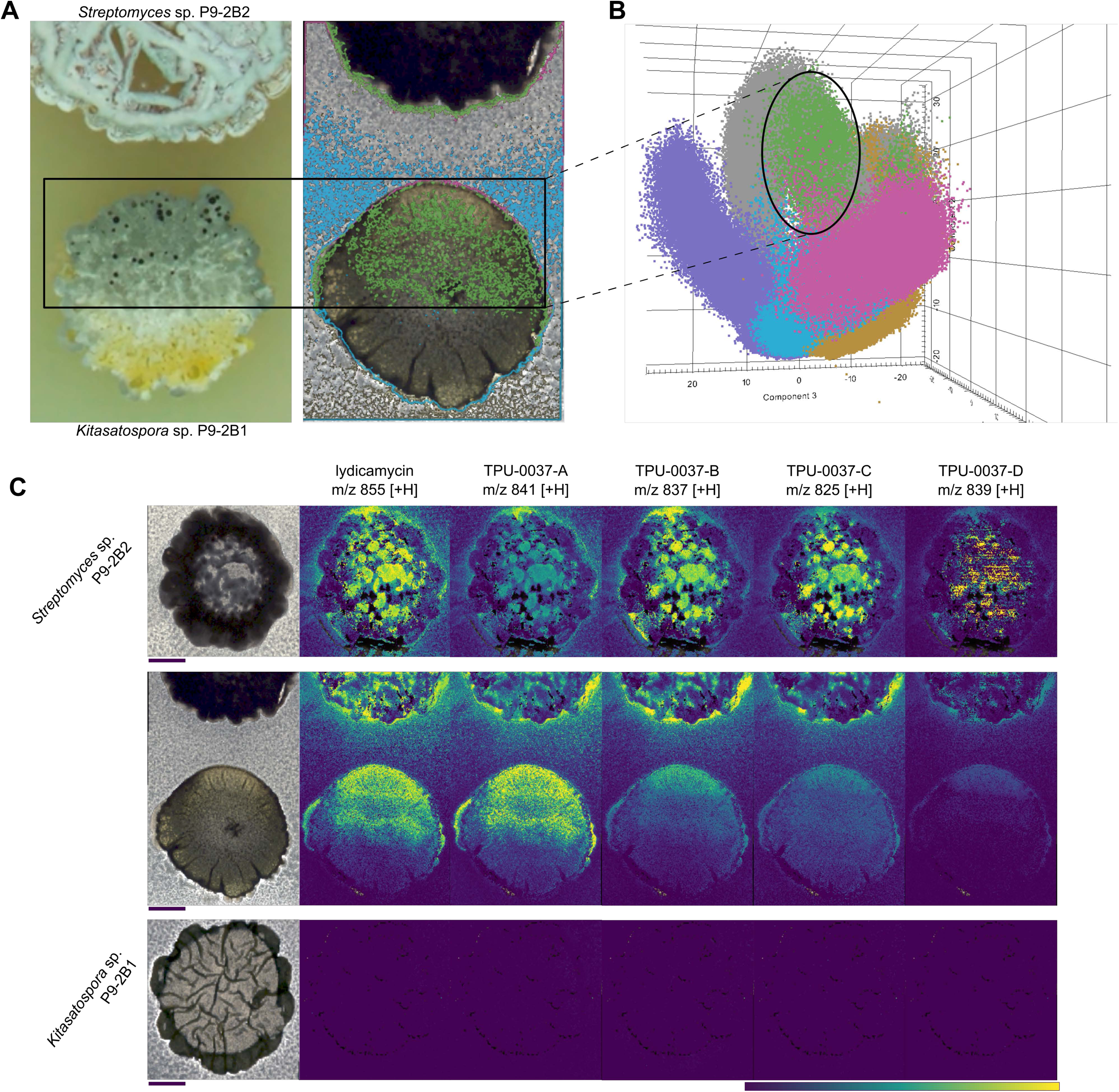
MALDI-MSI of *Kitasatospora* sp. P9-2B1 and *Streptomyces* sp. P9-2B2. (A) Digital microscope image of sporulation and interaction zone between the actinobacteria (left) and MSI sampling region with segmentation differentiating P9-2B1(blue), sporulation zone of P9-2B1 (green) and P9-2B2 (magenta). (B) 3D PCA plot of MSI spectra with two additional segments of the P9-2B1 monoculture (lavender) and P9-2B2 monoculture (orange). (C) Ion images of the various lydicamycins produced (TPU-0037-A through D are congeners of lydicamycin). Scale bars: 4mm. Colored scale bar indicates relative ion abundance with hotspot removal activated.

Using Bruker SciLS, we focused on spatially co-localized features ranging of *m/z* 800-860. MSI of the monocultures show these features are present in and around P9-2B2, the strain causing sporulation. These features were present in the sporulation area of P9-2B1, giving a strong indication these are the responsible features for the observed sporulation. Furthermore, Principal Component Analysis revealed the sporulation zone of P9-2B1 (Fig. 2B) was more similar to the surrounding agar and colony of P9-2B2 than of the surrounding agar and non-sporulated colony of P9-2B1. Additionally, the P9-2B1 monoculture was separated from the sporulation zone, indicating they share no metabolic overlap. Using BGC predictions from antiSMASH (26) and accurate mass comparison to NPAtlas(27) and MS/MS matching to the GNPS library, we hypothesized the responsible metabolites to be lydicamycins.

Lydicamycins are Non-Ribosomal Peptide Synthetase–Polyketide Synthase (NRPS-PKS) hybrids, recently named arginoketides(28), that have proposed biosynthesis starting from an enzymatically converted arginine(29) or a separate enzymatic process (30). Five known derivatives were originally described (lydicamycin and TPU-0037-A,B,C,D) and MSI was able to observe all known derivatives (Fig. 2C) (31, 32). P9-2B2 produces lydicamycins, which are diffused through the agar and then taken up by P9-2B1, principally in the sporulation zone. On top of the five known lydicamycins detected via MSI, additional lydicamycins were also detected that linked to the known lydicamycins via GNPS molecular networking (Fig. S6)). Three features (*m/z* 809.5084 [M+H]^+^, 823.5110 [M+H]^+^, and 853.5326 [M+H]^+^) are structurally related to TPU-0037-B based on molecular networking analysis (Fig. 3A), which contains an additional degree of unsaturation based on accurate mass formula calculations. One feature (*m/z* 827.5167 [M+H]^+^) is structurally related to TPU-0037-A and C. A recent study which also utilized molecular networking to identify new derivatives of lydicamycins also proposed these new congeners which we detect, including *m/z* 811.511 [M+H]^+^, which links in their study to TPU-0037-A (30). They further identify several other features which we also observe (*m/z* 843.5352, 857.5474, 869.53 and 871.54; [M+H]^+^) and additionally, we observe putative analogs (*m/z* 809.5084, 829.5238; [M+H]^+^) which they do not, pointing to the fact there are further products in the biosynthetic pathway to uncover (raw data available). Based on the accurate mass and matching fragmentation spectra to the GNPS library, all confirmed and putative lydicamycins were assigned to Level 2 and Level 3 confidence, respectively, based on Schymanski’s rules (33). All confirmed and putative lydicamycin MS^2^ spectra were deposited into the GNPS Spectral Library (Table S2).

**Fig. 3.**
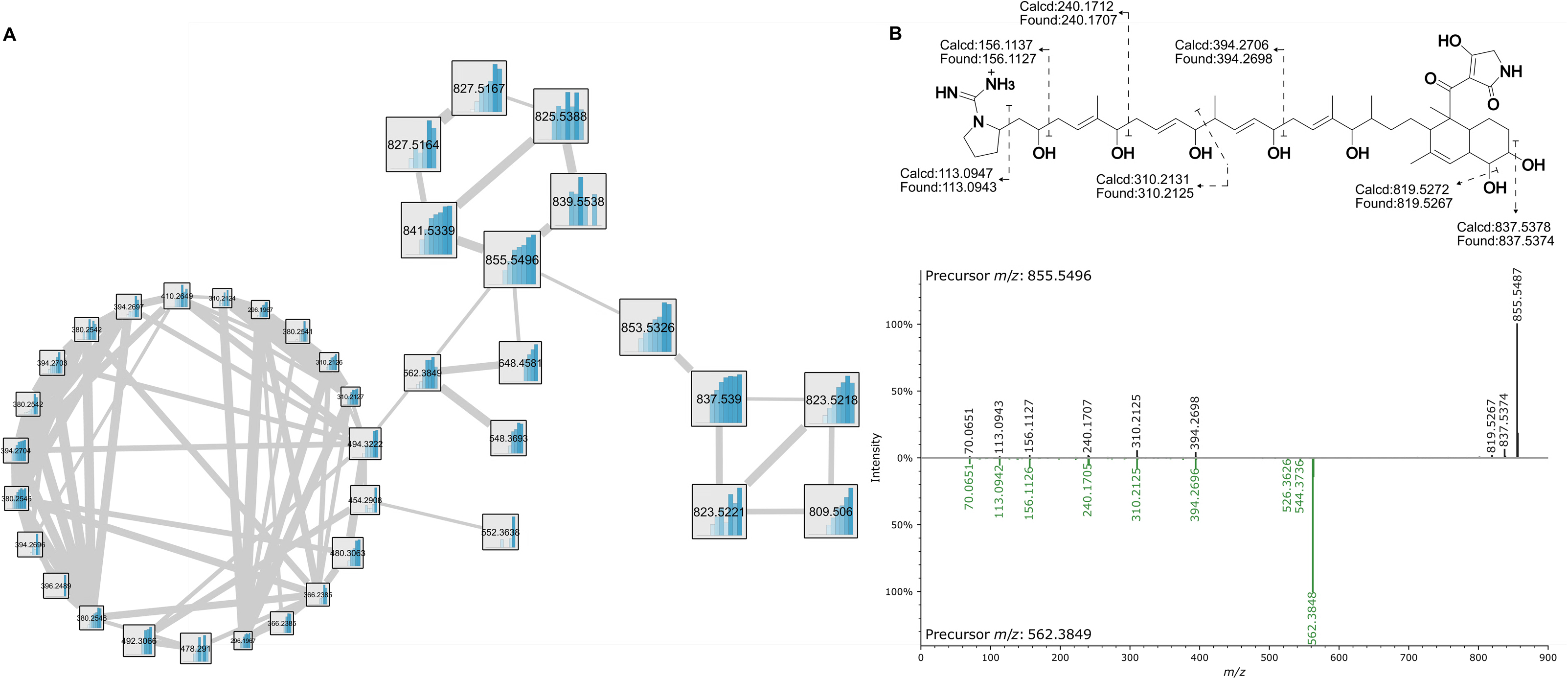
(A) Lydicamycin molecular family where increased edge width indicates higher fragmentation similarity. Internal bar graphs indicate peak area over each day from Day 1-10 (B) Fragmentation of lydicamycin (*m/z* 855.5496) and mirror plot of lydicamycin and the putative shunt product (*m/z* 562.3849). Green highlights indicate a cosine score-based match.

Lastly, an additional set of features was connected in the Feature Based Molecular Networking (FBMN) via lydicamycin (*m/z* 855) yet were smaller in *m/z* (Fig. 3A). Zhang et al. also observed these smaller features in their molecular network yet provided no explanation for their presence (30). Due to their connectivity, we knew their fragmentation was similar, pointing to their structural similarity as well (Fig. 3B). Upon further investigation, the fragments at *m/z* 394.2696, 310.2125, 240.1707, and 156.1126 can all be traced to the polyketide backbone (Figure 3B) and to the previously reported fragmentation (31), on the C-C bond adjacent to hydroxyls. Due to this fragmentation similarity, we hypothesize these remaining features are shunt products or byproducts of lydicamycin biosynthesis (35, 36). The study by Deng et al. (36) shows a similar metabolic flux to shunt production as we observe here; upon production of lydicamycins at day 4, we see immediate detection of potential shunt products and an increase towards days 9 and 10. Additional annotations from the GNPS library can be found in Fig. S7.

### Lydicamycin production is strongly tied to aerial mycelium production

Due to the role time plays in our coculture system, we sought out to investigate when lydicamycins are produced and attempt to link that to morphogenesis. We conducted an untargeted metabolomics analysis over 10 days, where each day a new plate containing three separate macro-colonies were imaged and extracted using plug extraction. Principal Coordinate Analysis (PCoA) in Qiime2 View(37) allowed us to observe the relative abundances of each feature over this 10-day time scale (Fig. 4A). Furthermore, Pearson r correlation distinguishes three sample groupings: days 1-3, 4-5 and 6-10 (Supplementary Fig. S8). Nodes from the FBMN (Fig. 4A) indicate all the lydicamycins and the putative shunt products increased in production up to day 9 (saturation of the MS). When comparing this data to the microscopy images taken on each of these days, we observe morphogenesis and the onset of sporulation strongly correlates with the three metabolomics-based sample groupings: Day 1-3 are defined by vegetative growth and no detectable lydicamycin production, days 4-5 are the onset of aerial mycelium and lydicamycin production and days 6-10 are defined by maturation and sporulation along with the highest level of production (Fig. 4B and C). Supplementary microscopy images can be found in Supplementary Figure S9.

**Fig. 4.**
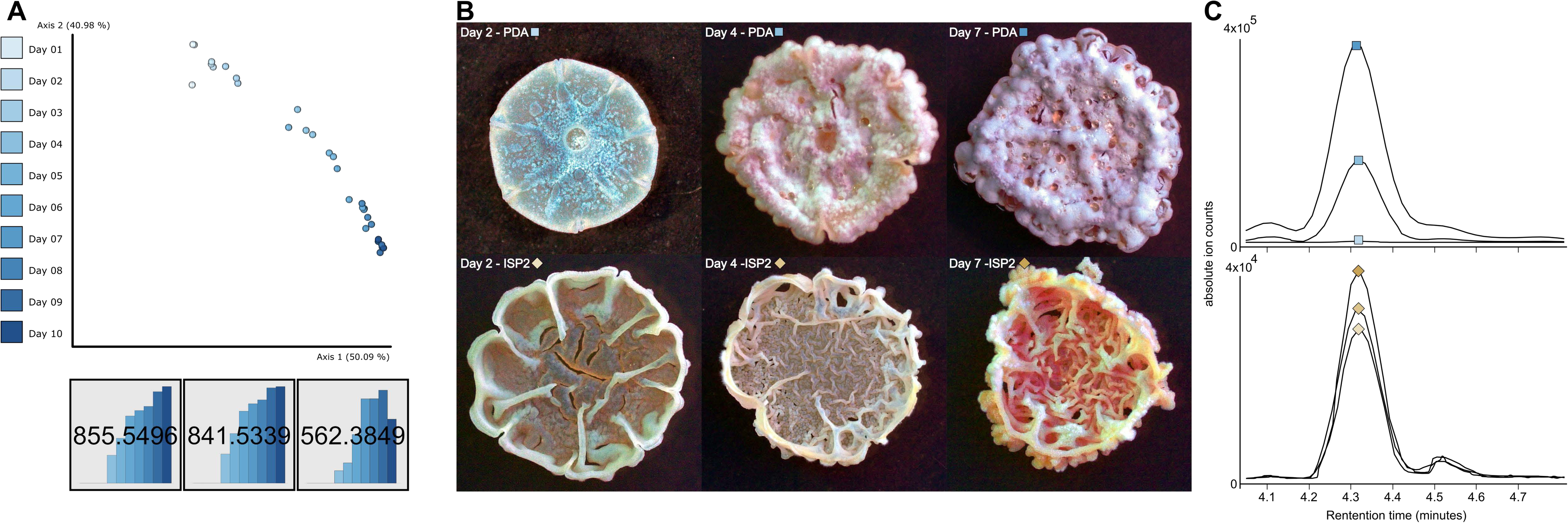
(A) PCoA showing the difference in metabolome over the cultivation time of P9-2B2 on PDA. Molecular networking nodes show the production of lydicamycins (*m/z* 855.5496 and 841.5339) and the putative shunt product, *m/z* 562.3849. (B) Microscopy of P9-2B2 colonies on PDA and ISP2 from 1, 4, and 7 days of growth on solid agar. (C) Extracted ion chromatograms of *m/z* 855.5496 corresponding to the timed images in B for P9-2B2 grown on PDA (top) and ISP2 (bottom).

In early experiments, we also observed when the two *Streptomyces* isolates were cocultivated on ISP2 agar, neither the receiver strain P9-2B1 had sporulation induced nor did P9-2B2 sporulate itself. Therefore, we hypothesized that lydicamycin production must be lacking due to both observations. The same untargeted time-based experimental setup as described above (with fewer timepoints) was carried out on ISP2 and the results showed no sporulation in P9-2B2 over 7 days (Fig. 4B). Surprisingly, lydicamycin production began on Day 1 when cultured on ISP2 and was stagnant through Day 7, but overall, the relative quantities were 10x lower than that on PDA. Based on the timing of lydicamycin production of PDA compared to that of ISP2 and their corresponding phenotypes, we concluded that the onset of P9-2B2 sporulation kickstarts lydicamycin production, a typical phenomenon of secondary metabolite production in *Streptomyces* (*38*).

### Lydicamycins induce morphogenesis in *Streptomyces coelicolor*

Having shown the effect of strain P9-2B2 and lydicamycin production on strain P9-2B1 in this study, we set out to explore if sporulation is also triggered in other environmentally relevant *Streptomyces* from our collection site. Two additional environmental actinobacteria (*Kitasatospora* sp. P9-2B3 and *Streptomyces* sp. P9-2B4) and *Streptomyces coelicolor* M145 and M1146 were cocultured with P9-2B2 and timelapse images for each were taken. Results showed that *Kitasatospora* sp. P9-2B3 and the two *S. coelicolor* isolates exhibited the same phenotype change as P9-2B1 (Fig. S10). Sporulation waves began to appear for P9-2B3 starting on Day 7 and progressing towards full sporulation on Day 10 (Fig. S10), similar to P9-2B1. Based on phylogenetic tree reconstruction, strains P9-2B1 and P9-2B3 are both highly similar to each other, and likewise for P9-2B2 and P9-2B4 (Figures S1-4). Timelapse videos of each interaction can be found in the supplementary data (Videos S1-4).

Sporulation of *Streptomyces coelicolor* M145 has been shown to be inhibited by prodiginine (39), which could explain why we observed a delayed sporulation on day 10 and did not fully sporulate by day 21. Strong prodiginine production could be seen starting on day 4, which has been implicated in delaying sporulation in *Streptomyces coelicolor* (39), and actinorhodin production was observed in different sporulating regions across days 10-21 (Video S6). Therefore, to remove the effect of the prodiginines, we carried out subsequent work on *Streptomyces coelicolor* M1146 which contains four deleted BGCs, including prodiginines (40). In contrast to M145, sporulation in M1146 occurred after 4 days, with full sporulation seen 24 hours later (Video S7).

To confirm our lydicamycin inducible-sporulation hypothesis based on imaging, metabolomics, and genomics, we generated a *lyd*-deficient mutant to test in coculture. The first core polyketide synthase gene *lyd60* was targeted and inactivated by converting a TGG (Trp) codon at position 57 into the stop codon TAA using CRISPR-based base-editing tool CRISPR-BEST (16). The deficient mutant (*lyd*60^STOP^) was confirmed by Sanger sequencing of the editing site, no detectable lydicamycin after 7 days of growth on PDA using LC-MS (Figure S11), and with cocultivations showing a lack of induced morphogenesis in *Kitasatospora* sp. P9-2B1 (Fig 5). Subsequent timelapse images were taken over 10 days and no sporulation was observed in the receiver strain, P9-2B1 (Video S8). *lyd6*0^STOP^ had observably different growth in the first two days from the wild type (WT). The first indication of aerial mycelium in WT P9-2B2 appears on Day 1 Hour 13 compared to Day 4 for the *lyd*-deficient mutant (Videos S9-10), indicating the inactivation of the *lyd* BGC clearly delays development in P9-2B2. To further confirm lydicamycin’s role in inducible morphogenesis, pure lydicamycin (*m/z* 855) was added into an agar well and induced sporulation in P9-2B1 but only at 100 µg and a longer interval (9 days) than when cocultured (6-7 days). Sporulation assays were conducted on pregrown cultures for 3 days, otherwise, P9-2B1 was inhibited by pure lydicamycin when cultured simultaneously, indicating its overall antimicrobial effect.

**Fig. 5.**
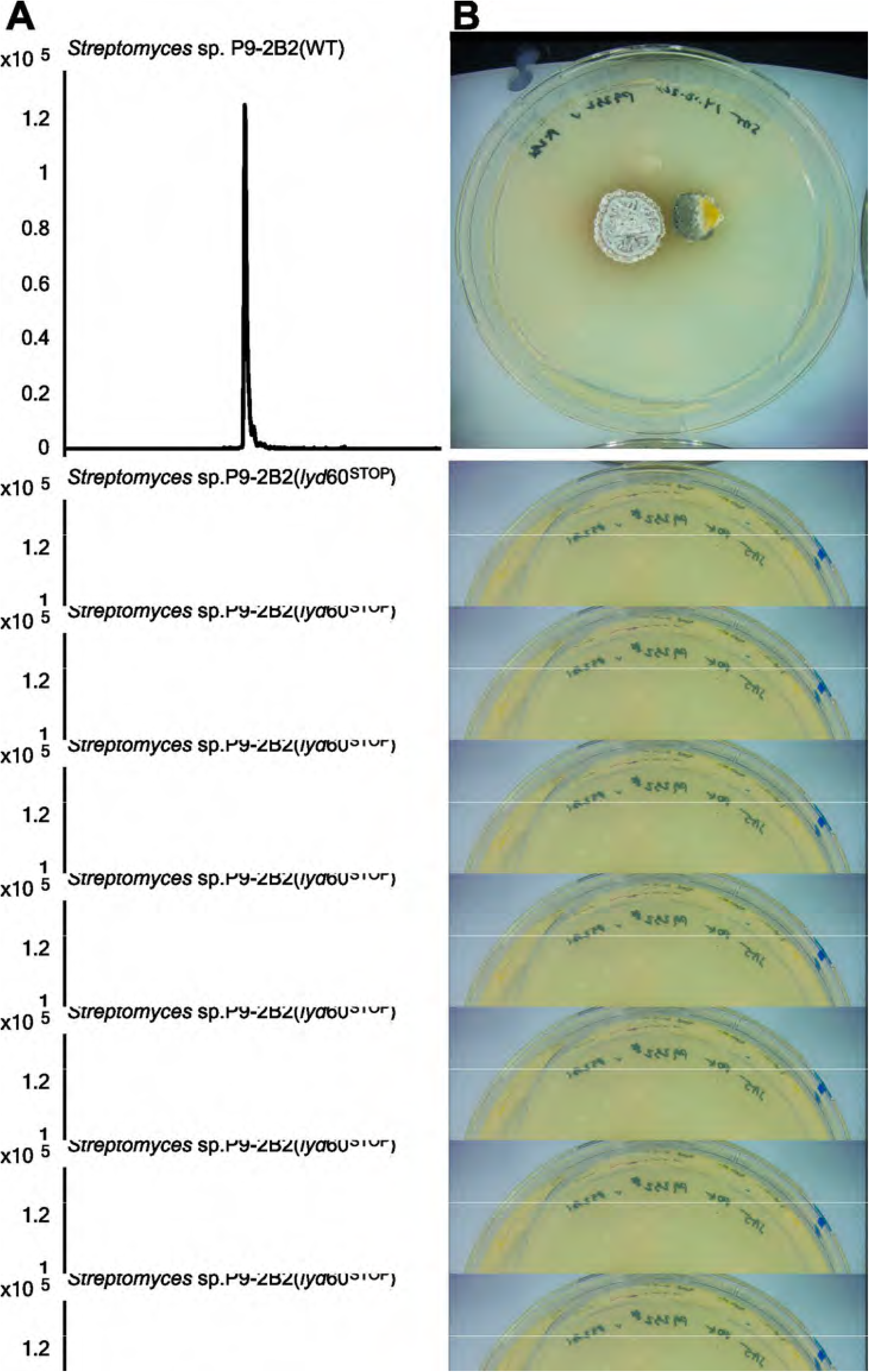
(A) Extracted ion chromatogram of lydicamycin (*m/z* 855.5496) in co-cultivation of WT P9-2B2 and P9-2B1 (top), cocultivation of *lyd*60*^STOP^*and P9-2B1 (middle) and monoculture of P9-2B1 (bottom). (B) Corresponding images from each cocultivation where P9-2B2 is present on the left and P9-2B1 on the right.

### Genes controlling aerial mycelium are expressed upon increased lydicamycin exposure

Morphogenesis is a phenomenon brought on in Streptomycetes by nutrient depletion and is irreversible (38). Based on what we have observed in this study, it is clear that lydicamycins produce a similarly stressful environment for the receiver strains unrelated to nutrient availability, therefore, we wanted to evaluate the transcriptional changes ongoing during increasing lydicamycin exposure. We evaluated the transcriptome of the receiver strain (M1146) in monoculture (MC) and coculture (CC) over four timepoints (2, 4, 7, and 9 days). In this sampling day 2 represents a datapoint where no lydicamycins are detectable (by LC-MS), where lydicamycins are first observed in LC-MS data (day 4), when exposure to high concentrations of lydicamycin are present (day 7), and day 9 where aerial mycelium begins to appear (and when we expect spores to form quickly thereafter). On day 2, there were no differential genes observed between CC and MC. Day 4 (Fig. 6) and day 9 (Fig. 7) both show extensive differential expression (log_2_ and p_adj_ < 0.01), 123 and 410 total genes, respectively, encompassing development, cell envelope stress, sigma factors and secondary metabolism. Day 7 surprisingly showed very little differential expression between MC and CC. Data for days 4 and 9 can be found in Supplementary file 2 and 3, respectively.

**Fig. 6.**
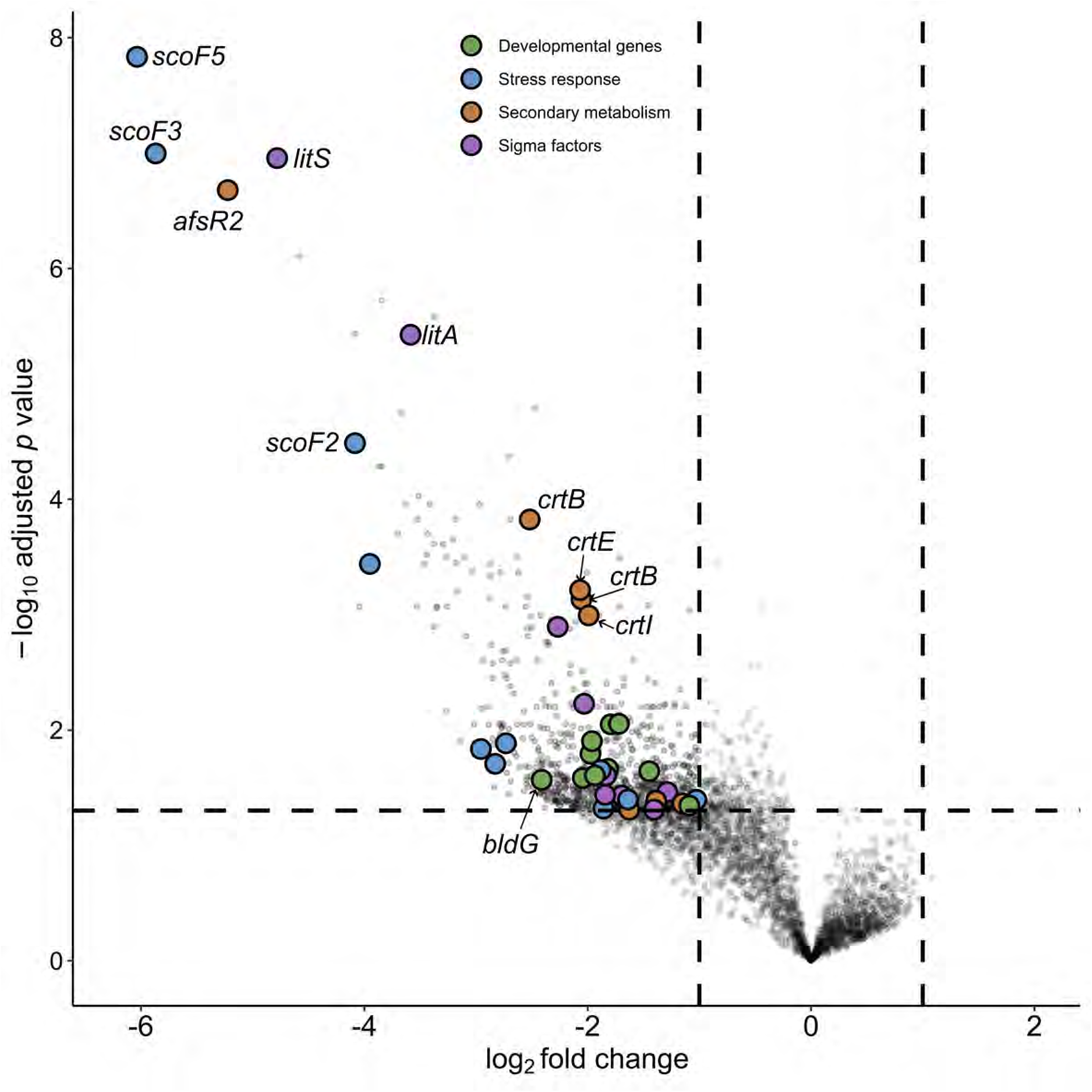
Volcano plot showing differentially expressed genes in *Streptomyces coelicolor* M1146 between day 4 monoculture vs. coculture. Genes with a negative log_2_ fold change were relatively more abundant in coculture (*n* = 4) and genes with a positive log_2_ fold change were relatively more abundant in monoculture (*n* = 4). Key genes associated to development, sigma factors, stress and secondary metabolism were annotated using SCO identifiers via StrepDB (<u>StrepDB</u> (streptomyces.org.uk).

**Fig. 7.**
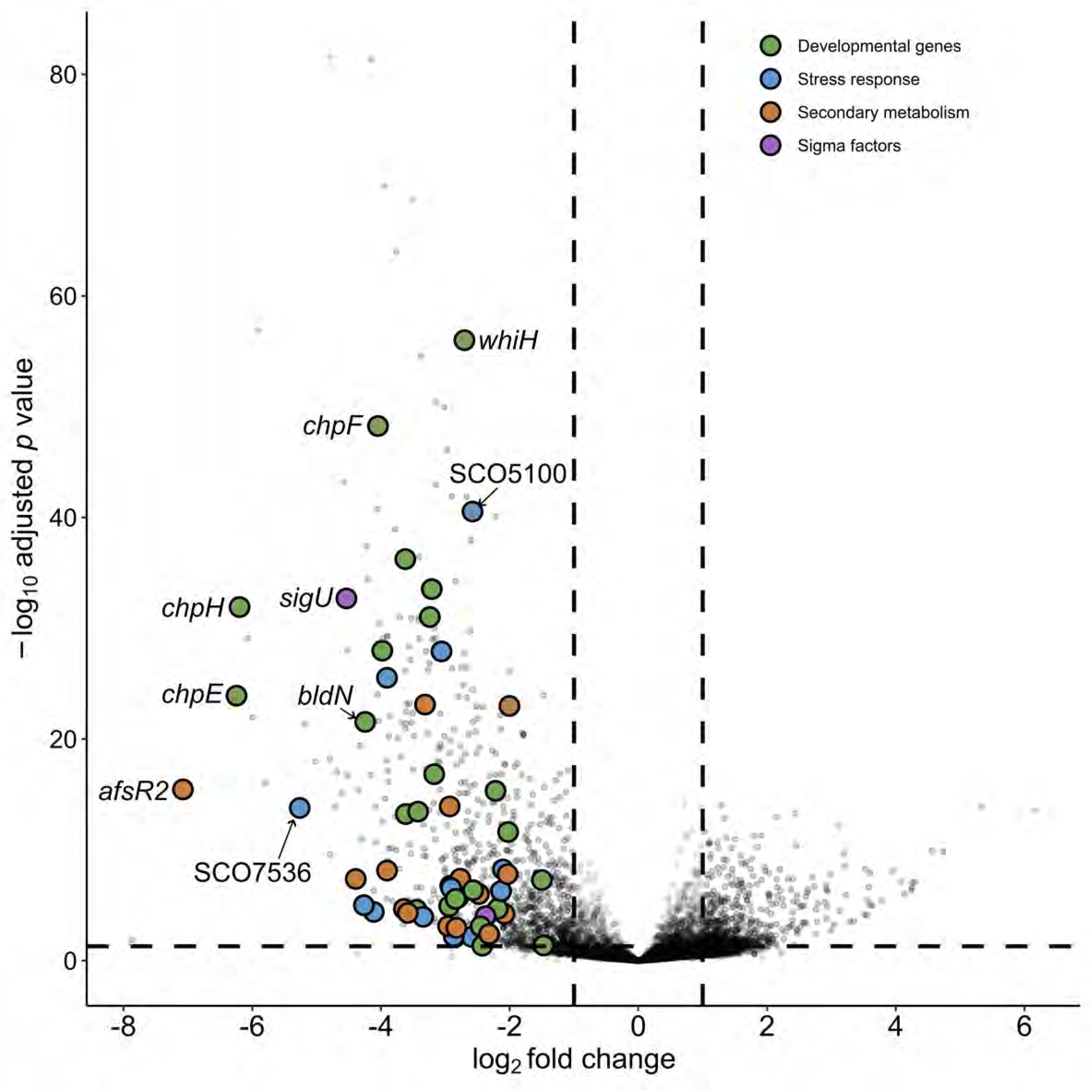
Volcano plot showing differentially expressed genes in *Streptomyces coelicolor* M1146 between day 9 monoculture vs. coculture. Genes with a negative log_2_ fold change were relatively more abundant in coculture (*n* = 4) and genes with a positive log_2_ fold change were relatively more abundant in monoculture (*n* = 4). Key genes associated with development, sigma factors, stress and secondary metabolism were annotated using SCO identifiers via StrepDB (<u>StrepDB</u> (streptomyces.org.uk).

Compared to the *Streptomyces coelicolor* M1146 MC, *bldD* (SCO1489) and *bldG* (SCO3549) is differentially expressed in day 4 CC and we further observe the early chaplin genes (*chpE* and *chpH*) along with *sapA* (SCO4049) differentially expressed, to produce the hydrophobic layer allowing spores to form later (Fig. 6). Multiple sigma factors ( *σ^H^,σ^I^, σ^N^, σ^8^*, SCO4005, SCO3613) and regulators (*wblA*, *afsQ1/Q2*, *afsR2*, *phoR*, SCO5147, and *cvnA2/B2*) which are responsible for differentiation and antibiotic production(41) were also differentially expressed on day 4 CC (log2 foldchange < −2). We also observe conservon loci (*cvnA1/B1/D1* and *cvnA13*) being differentially expressed at day 4 CC. These two loci are co-regulated with *cvn8* which has been shown to directly stimulate antibiotic production during interspecies interactions (42).

Importantly, we observed differential expression of genes associated with various stress responses in M1146 (Fig. 5): oxidative stress response (*nur*, σ^R^, σ^8^, SCO5749, *osaB*, *osaC*), osmotic stress (σ^H^ and σ^I^), and especially shock proteins (HspR, ScoF and ScoF1-5), all of which are general responses to cell envelope stress (43). We also see differential expression of the carrotenoid BGC *crt* and the neighboring sigma factor (σ^LitS^) that is both light and stress induced (44). BlastP on *crtY* shows a 59% homology to a putative lycopene cyclase (WJY42522.1) in *Streptomyces* sp. P9-2B2, therefore it is logical that the yellow pigmentation in our original observation (Fig. 1A) that preceded aerial mycelium is also a carotenoid derived from this BGC.

### Lydicamycins elicit a transcriptional response similar to cell wall targeting antibiotics

On day 9 CC, where we observed half of the M1146 colony had transitioned to aerial hyphae, genes tightly associated to aerial hyphae formation (*bldN*, *bldM*, *wblA*) along with hydrophobic proteins and peptides (HpB, ChpD-H, SapA-B) were differentially expressed (Fig. 7) (41). Furthermore, we see differentially expressed genes associated with kickstarting sporulation (*ssgA*, *ssgB*, *ssgR*, s*poIIE*) and conversion of aerial hyphae to spores (*whiH*), all of which are activated by the global regulator *bldD* (log2 foldchange < −2) (41). *wblH* (log2 foldchange = - 2.56), which is controlled via *whiA*(*45*), showed a similar transcriptional pattern to *wblA* and *bldN* and may have a similar role but further investigations are required.

Day 9 is not only defined by conversion of aerial mycelium and the beginning of sporulation, but we also observe further signs of cell envelope stress (Fig. S12). We see differential expression of a hypothetical protein (SCO5100) homologous to *ytrA* from *Bacillus subtilis*, a GntR family repressor involved in cell envelope stress response to cell wall targeting antibiotics (46). *σ^E^*is induced by a wide range of cell wall targeting antibiotics(47) and is a major marker of cell envelope stress in *Streptomyces coelicolor* (48). Along with a σ^E^-like protein (SCO4005), we observe differential expression of 12/28 tightly associated *σ^E^* regulons seen by Pospíšil et al. when *S. coelicolor* was exposed to ethanol stress (49). We also see differential expression of *σ^U^* (Fig. S7), which modulates functional proteins directly associated to counteracting cell envelope stress as well as modulating morphological differentiation(50), and several additional sigma factors (*σ*^8^, σ^F^, σ^H^, σ^I^, σ^N^ and σ^R^). Additional evidence of cell wall stress includes differential expression of SCO6091 and SCO7536, both homologs of Mycobacterial membrane protein Large (MmpL) transporters which are pumps that send lipids to the cell envelope under stress (51). In terms of secondary metabolism, we observed differential expression of genes associated to coelibactin biosynthesis, however it is unclear whether this is due to a generic stress response(52) or coelibactin’s ecological role in sporulation (53). We further observed differential expressions of two lanthipeptide precursor peptides (SCO6931 and SCO6932) that recently have been shown to be a part of anti-phage defense (54). Combined with the observed differential expression of a putative abortive phage infection resistance protein (FQ762_32655) that trigger cell suicide to prevent phage propagation (55), lydicamycin-induced morphogenesis or morphogenesis in general may trigger anti-phage defenses as part of cell stress response.

## Discussion

The induction of sporulation by lydicamycin makes this one of a handful of described SMs with the same ecology. The two other SMs described with sporulation inducing effects are goadsporin and desferrioxamine E, with our results more similarly mirroring goadsporin as both are antibacterial and sporulation inducing(9). Yamanaka et al. reported the sporulation-inducing activity of desferrioxamine E and confirmed the observation via BGC knockout studies (56). They showed their susceptible strain, *S. tanashiensis*, lacked the *des* BGC and the siderophore transporter, therefore making it prone to iron starvation. However, this is most likely a rare case amongst *Streptomycetes* as the majority of genomes (71 %) encode for the highly conserved desferrioxamine BGC (57). Additionally, there are several metabolites further responsible for differentiation, specifically germination, which are described (38), pointing to the fact that *Streptomyces* SMs may play larger roles in its life cycle than previously anticipated.

According to us, there are two main factors leading to the underreported number of sporulation inducing SMs: (1) the low number of pathogenic *Streptomycetes* discourages biological testing in traditional drug discovery efforts and (2) low compound yields make it difficult to test against a broad enough range of bacteria. The induction of morphological differentiation is a viable strategy for increasing the production of antibiotics (58, 59) and may be a route for eliciting cryptic biosynthetic pathways through subinhibitory concentrations, yet the latter requires further investigation. Nevertheless, we require ecological studies to identify metabolites capable of inducing sporulation to begin investigating the potential downstream effects. Furthermore, as we reveal further ecological functions of SMs, the ability to utilize microbes as biocontrol agents will also gain further attention, beyond their high interest already garnered.

Traditional analytical techniques like HPLC or LC-MS lack the ability to quickly discern metabolites which may be involved in cocultures (unless they are produced in high titres), either lacking noticeable differences in UV chromatograms or difficulty deconvoluting many features, respectively. On the other hand, MSI is a burgeoning technique equipped uniquely to handle these types of samples, as seen through this study and others (10, 60, 61). Using MSI, we were able to quickly define potential candidate features based on their spatial distribution and through dereplication via MS and genome mining, propose and confirm the sporulation-inducing nature of lydicamycins. MSI suffers from topological challenges and actinobacteria and other filamentous microbes can be difficult to chemically image, therefore, modified strategies like what we have demonstrated or new techniques may be required to increase the utility of MSI. Furthermore, MS-based metabolomics like molecular networking and multivariate statistics continue to be able to differentiate microbial metabolomes, as we have demonstrated with our temporal study. While transcriptomics and proteomics have traditionally been utilized to track changes over time, metabolomics offers visibility towards the end of metabolic processes. Ultimately, all three -omics provide separate details in the life of a microbe and integration of these data will begin to reveal the larger picture of SM production.

We have shown that via MSI, the characterization of lydicamycins as the responsible metabolites in a dual *Streptomyces* coculture is the optimal technique for these agar-based setups. Not only were the standard congeners detected but several additional newly discovered and putative derivatives were also visible in the data. Temporal production of the lydicamycins nicely correlated with self-induced sporulation of the producing strain and the generation of a deficient mutant significantly delayed the onset of sporulation when compared to the wild type. Lastly, through transcriptomics, we observed the morphogenesis transitions of M1146 alongside increasing lydicamycin exposure and observed several differentially expressed genes pertaining to cell envelope stress. This study provides an important primer for tracking down the responsible metabolites in microbial interactions but also begins to illuminate the ecological role and temporal nature of lydicamycins.

## Acknowledgements

The study was supported by the Danish National Research Foundation (DNRF137) for funding the Center for Microbial Secondary metabolites (CeMiSt). We would also like to acknowledge funding from Novo Nordisk Foundation (grant NNF19OC0055625) for the infrastructure “Imaging microbial language in biocontrol (IMLiB). Z.Y. acknowledges funding from the China Scholarship Council (202004910340). TW would like to acknowledge funding from The Novo Nordisk foundation (grant NNF20CC0035580) for the Center for Biosustainability. Finally, we acknowledge the DTU Metabolomics Core for usage of LCMS equipment as well as daily maintenance.

## Data Availability

LC-MS data can be found in GNPS-MassIVE at MSV000092216 and MSI data can be found at Metaspace at https://metaspace2020.eu/project/jarmusch-2023. Isolate sequencing data have been deposited at the NCBI BioProject database with accession number PRJNA985726. FBMN workflow can be found here: https://gnps.ucsd.edu/ProteoSAFe/status.jsp?task=a3dd5a7a459e475d8ca492ef7d05f3b7. Raw RNA-seq data have been deposited at NCBI SRA with accession number PRJNA1123431. All code used in this study can be found at https://figshare.com/articles/software/Files_for_the_analysis/26038807 and the output from the kallisto analysis can be found at https://figshare.com/articles/dataset/Input_files_for_DeSeq2_analysis/26029459.

