## Supplementary Data for "Lydicamycins Induce Morphological Differentiation in Actinobacterial Interactions"

**DSMZ Type (Strain) Genome Server**

The analysis also made use of recently introduced methodological updates and features^1^. Information on nomenclature, synonymy and associated taxonomic literature was provided by TYGS's sister database, the List of Prokaryotic names with Standing in Nomenclature (LPSN, available at https://lpsn.dsmz.de)^1^. The results were provided by the TYGS on 2024-11-20. The TYGS analysis was subdivided into the following steps:

*Determination of closely related type strains*

Determination of closest type strain genomes was done in two complementary ways: First, all user genomes were compared against all type strain genomes available in the TYGS database via the MASH algorithm, a fast approximation of intergenomic relatedness^2^, and, the ten type strains with the smallest MASH distances chosen per user genome. Second, an additional set of ten closely related type strains was determined via the 16S rDNA gene sequences. These were extracted from the user genomes using RNAmmer^3^ and each sequence was subsequently BLASTed^4^ against the 16S rDNA gene sequence of each of the currently 22086 type strains available in the TYGS database. This was used as a proxy to find the best 50 matching type strains (according to the bitscore) for each user genome and to subsequently calculate precise distances using the Genome BLAST Distance Phylogeny approach (GBDP) under the algorithm 'coverage' and distance formula d5^5^. These distances were finally used to determine the 10 closest type strain genomes for each of the user genomes.

*Pairwise comparison of genome sequences*

For the phylogenomic inference, all pairwise comparisons among the set of genomes were conducted using GBDP and accurate intergenomic distances inferred under the algorithm 'trimming' and distance formula d5^5^. 100 distance replicates were calculated each. Digital DDH values and confidence intervals were calculated using the recommended settings of the GGDC 4.0^1,5^.

*Phylogenetic inference*

The resulting intergenomic distances were used to infer a balanced minimum evolution tree with branch support via FASTME 2.1.6.1 including SPR postprocessing^6^. Branch support was inferred from 100 pseudo-bootstrap replicates each. The trees were rooted at the midpoint^7^ and visualized with PhyD3^9^.

**Inactivation of the lydicamycin BGC**

All primers used were synthesized by IDT (Integrated DNA Technologies, USA) and are listed in Supplementary Table S1. Plasmids and genomic DNA purification, polymerase chain reaction (PCR), and cloning were conducted according to standard procedures using manufacturer protocols. PCR was performed using Q5® High-Fidelity 2X Master Mix (New England Biolabs, USA). DNA assembly was done by using NEBuilder HiFi DNA Assembly Master Mix (New England Biolabs, USA). DNA digestion was performed with FastDigest restriction enzymes (Thermo Fisher Scientific, USA). NucleoSpin Gel and PCR Clean-up Kits (Macherey-Nagel, Germany) were used for DNA clean-up from PCR products and agarose gel extracts. One Shot Mach1 T1 Phage-Resistant Chemically Competent *E. coli* (Thermo Fisher Scientific, USA) was used for cloning. NucleoSpin Plasmid EasyPure Kit (Macherey-Nagel, Germany) was used for plasmid preparation. Sanger sequencing was carried out using a Mix2Seq Kit (Eurofins Scientific, Luxembourg). All DNA manipulation experiments were conducted according to standard procedures using manufacturer protocols.

**RNA-seq analysis**

Raw RNA reads (average 1.41 107 +- 2.03 106 SD reads per sample) were trimmed and filtered (average 1.41 107 +- 2.03 106 SD reads per sample) with fastp^1^ using default settings, and SSU rRNA was removed using SortMeRNA(v4.3.6)^2^ with the Silva database (smr_4.3_default_db.fasta)^3^ prior gene expression analysis (average 1.21 107 +- 1.84 106 SD reads per sample). Genome of Streptomyces coelicolor A3(2) (GCA_008931305.1) was retrieved from NCBI GenBank and converted into a fasta-formatted transcript file using gffread (v0.12.7).^4^ For transcript abundances Kallisto (0.46.2)^5^ was used to pseudo align the reads against the fasta-formatted transcript file with 100 bootstrap replicates and default settings. More than 88 % on average of the reads could be pseudoaligned. Differential expression analysis was performed with DESeq2 (v1.42.1)^6^ in Rstudio (v2023.12.1)^7^ running R (v4.3.3) with the Tidyverse framework (2.0.0)^8^. In order to retrieve the SCO identifcation numbers, we blasted the genes from the newly annotated genome (GCA_008931305.1) against the original Stretomyces coelicolor A2(3) (GCA_000203835.1) using Bowtie2^9^ and Samtools (v1.10)^10^ was used to sort and filter the results.
All of the used command-line and r-code for the complete analysis can be found on Figshare (<https://figshare.com/account/home#/projects/209641>).

1.	Chen, S., Zhou, Y., Chen, Y. & Gu, J. fastp: an ultra-fast all-in-one FASTQ preprocessor. Bioinformatics 34, i884–i890 (2018).

2.	Kopylova, E., Noé, L. & Touzet, H. SortMeRNA: fast and accurate filtering of ribosomal RNAs in metatranscriptomic data. Bioinformatics 28, 3211–3217 (2012).

3.	Quast, C. et al. The SILVA ribosomal RNA gene database project: improved data processing and web-based tools. Nucleic Acids Res. 41, D590–6 (2013).

4.	Pertea, G. & Pertea, M. GFF Utilities: GffRead and GffCompare. F1000Res. 9, (2020).

| Table S1. All bacteria, plasmid and primers used in this study | | |
| --- | --- | --- |
| **Strains** | **Description** | **Source/[Ref]** |
| One Shot™ Mach1™ T1 Phage-Resistant Chemically Competent *E. coli* | For routine plasmids maintenance and cloning | Thermo Fisher Scientific |
| *E. coli* ET12567/pUZ8002 | For conjugating plasmids into *Streptomyces* | Kieser et al. 2000^1^ |
| *Streptomyces* sp. P9-2B1 | Wild-type strain | In this work |
| *Streptomyces* sp. P9-2B2 | Wild-type strain | In this work |
| *Streptomyces* sp. P9-2B2/*lyd60* | *lyd60* inactivation mutant strain | In this work |
| *Streptomyces* sp. P9-2B4 | Wild-type strain | In this work |
| **Plasmids** |  |  |
| pCRISPR-cBEST | For C to T base editing | Tong et al. |
| pCRISPR-cBEST/*lyd60* | Modified plasmid for inactivation of *lyd*60 | In this work |
| **Primers** |  |  |
| pCRISPR-cBEST-*lyd60* | CCGGTTGGTAGGATCGACGGatcccacagttcctccggcgGTTTTAGAGCTAGAAATAGC | Oligo for protospacer sequence construction |
| ID-*lyd60*-F | ccgtagtcgtggtacatcac | Identification primer for mutant pCRISPR-cBEST/D*lyd60* |
| ID-*lyd60*-R | gaaagagtccgagccgatc | Identification primer for mutant pCRISPR-cBEST/D*lyd60* |

1.	Kieser, T., Bibb, M. J., Buttner, M. J., Chater, K. F. & Hopwood, D. A. Practical Streptomyces Genetics (The John Innes Foundation, 2000).

| Table S2. GNPS library annotations of lydicamycins | | | |
| --- | --- | --- | --- |
| CCMS library ID | Metabolite name | Adduct | m/z |
| [CCMSLIB00011430331](https://gnps.ucsd.edu/ProteoSAFe/gnpslibraryspectrum.jsp?SpectrumID=CCMSLIB00011430331#%7B%7D) | lydicamycin | M+H | 855.55 |
| [CCMSLIB00011430332](https://gnps.ucsd.edu/ProteoSAFe/gnpslibraryspectrum.jsp?SpectrumID=CCMSLIB00011430332#%7B%7D) | lydicamycin derivative: TPU-0037A | M+H | 841.53 |
| [CCMSLIB00011430333](https://gnps.ucsd.edu/ProteoSAFe/gnpslibraryspectrum.jsp?SpectrumID=CCMSLIB00011430333#%7B%7D) | lydicamycin derivative: TPU-0037D | M+H | 839.55 |
| [CCMSLIB00011430334](https://gnps.ucsd.edu/ProteoSAFe/gnpslibraryspectrum.jsp?SpectrumID=CCMSLIB00011430334#%7B%7D) | lydicamycin derivative: TPU-0037B | M+H | 837.54 |
| [CCMSLIB00011430335](https://gnps.ucsd.edu/ProteoSAFe/gnpslibraryspectrum.jsp?SpectrumID=CCMSLIB00011430335#%7B%7D) | lydicamycin derivative: TPU-0037C | M+H | 825.54 |
| [CCMSLIB00011430336](https://gnps.ucsd.edu/ProteoSAFe/gnpslibraryspectrum.jsp?SpectrumID=CCMSLIB00011430336#%7B%7D) | putative lydicamycin 1 | M+H | 809.51 |
| [CCMSLIB00011430337](https://gnps.ucsd.edu/ProteoSAFe/gnpslibraryspectrum.jsp?SpectrumID=CCMSLIB00011430337#%7B%7D) | putative lydicamycin 2 | M+H | 811.52 |
| [CCMSLIB00011430338](https://gnps.ucsd.edu/ProteoSAFe/gnpslibraryspectrum.jsp?SpectrumID=CCMSLIB00011430338#%7B%7D) | putative lydicamycin 3 | M+H | 823.52 |
| [CCMSLIB00011430339](https://gnps.ucsd.edu/ProteoSAFe/gnpslibraryspectrum.jsp?SpectrumID=CCMSLIB00011430339#%7B%7D) | putative lydicamycin 4 | M+H | 853.53 |
| [CCMSLIB00011430340](https://gnps.ucsd.edu/ProteoSAFe/gnpslibraryspectrum.jsp?SpectrumID=CCMSLIB00011430340#%7B%7D) | putative lydicamycin 5 | M+H | 869.53 |
| [CCMSLIB00011430341](https://gnps.ucsd.edu/ProteoSAFe/gnpslibraryspectrum.jsp?SpectrumID=CCMSLIB00011430341#%7B%7D) | putative lydicamycin 6 | M+H | 871.54 |

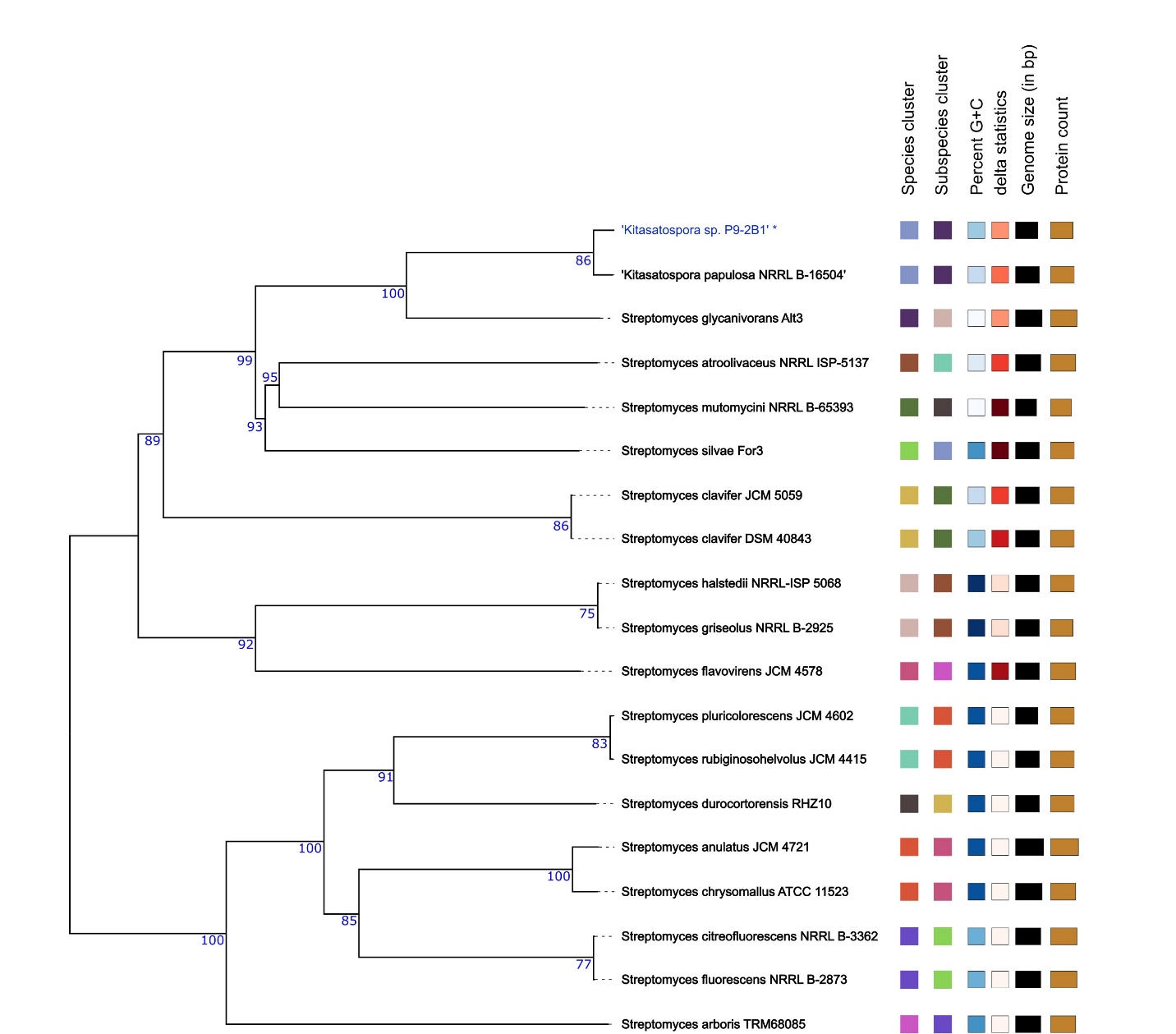

Figure S1. Phylogenetic tree of *Kitasatospora* sp. P9-2B1 from the DSMZ TYGS.

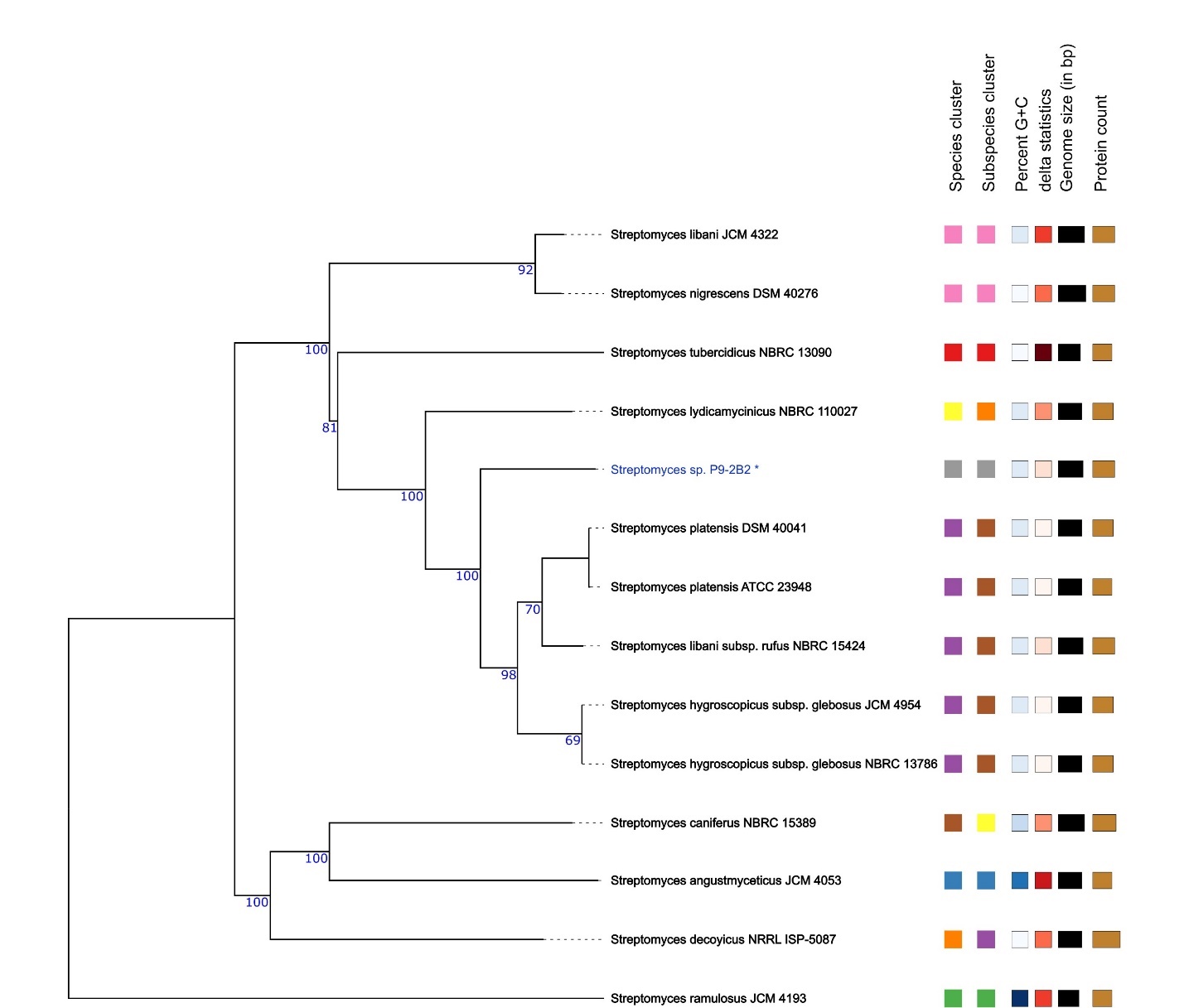

Figure S2. Phylogenetic tree of *Streptomyces* sp. P9-2B2 from the DSMZ TYGS.

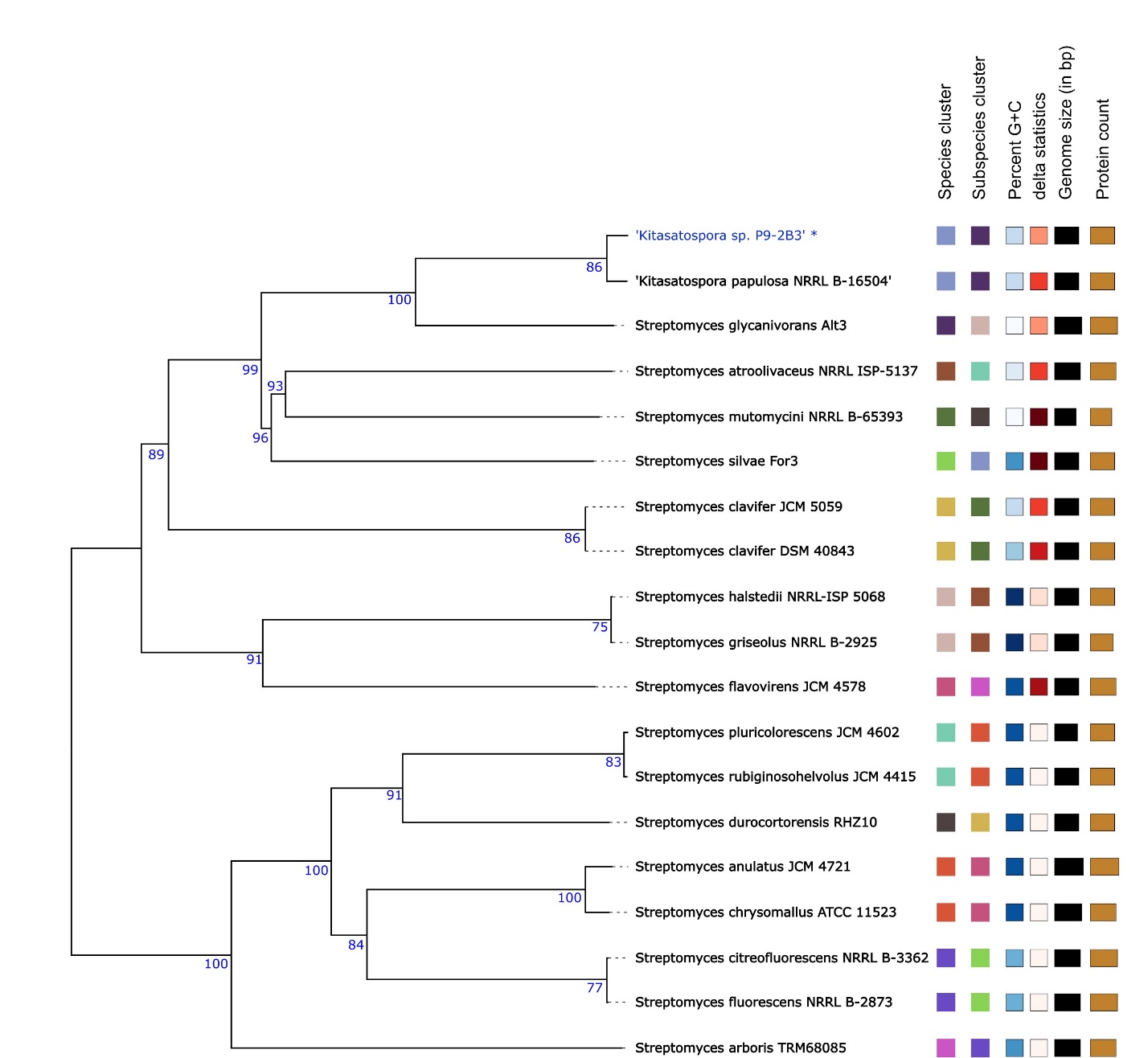

Figure S3. Phylogenetic tree of *Kitasatospora* sp. P9-2B3 from the DSMZ TYGS.

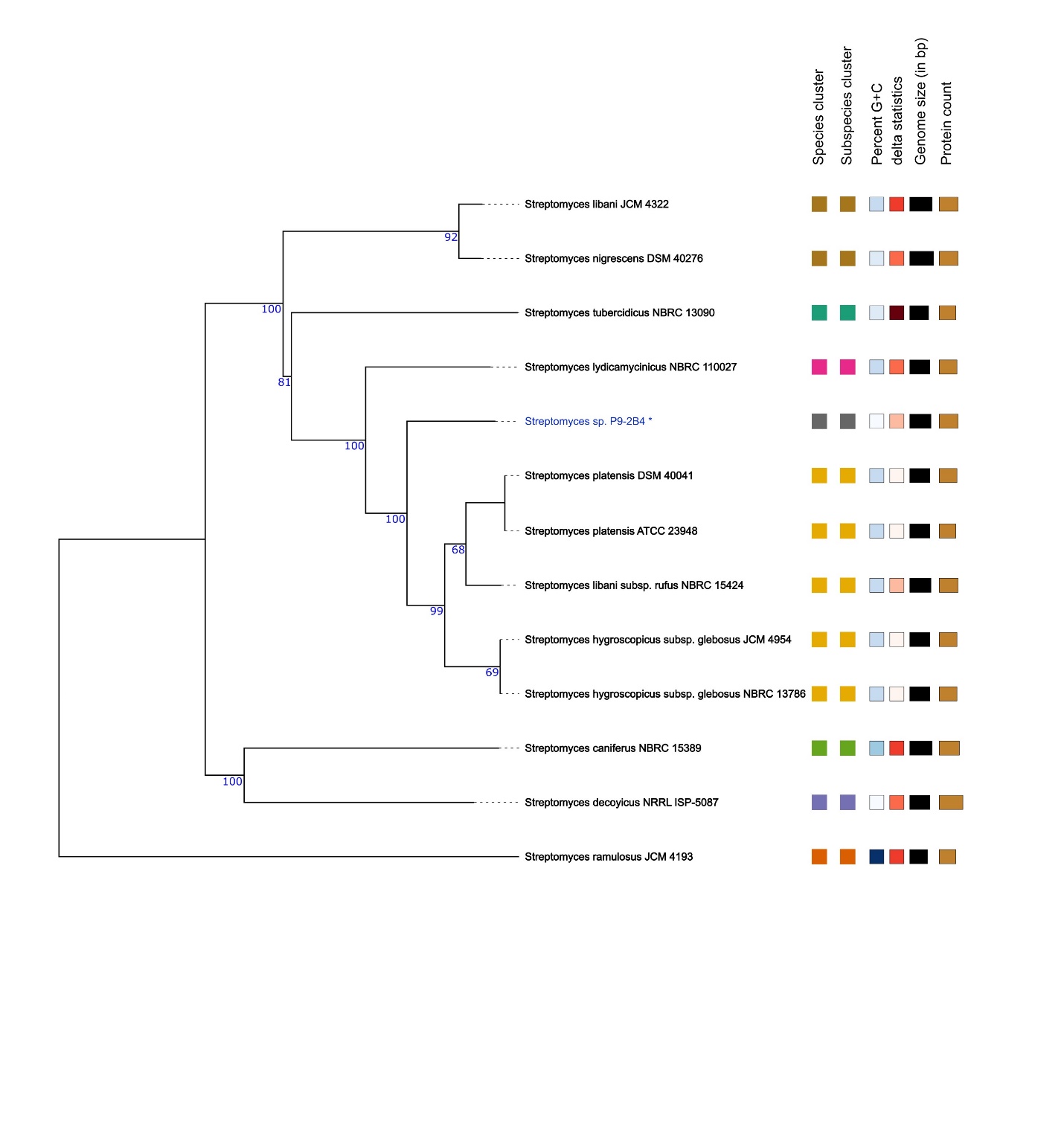

Figure S4. Phylogenetic tree of *Streptomyces* sp. P9-2B4 from the DSMZ TYGS.

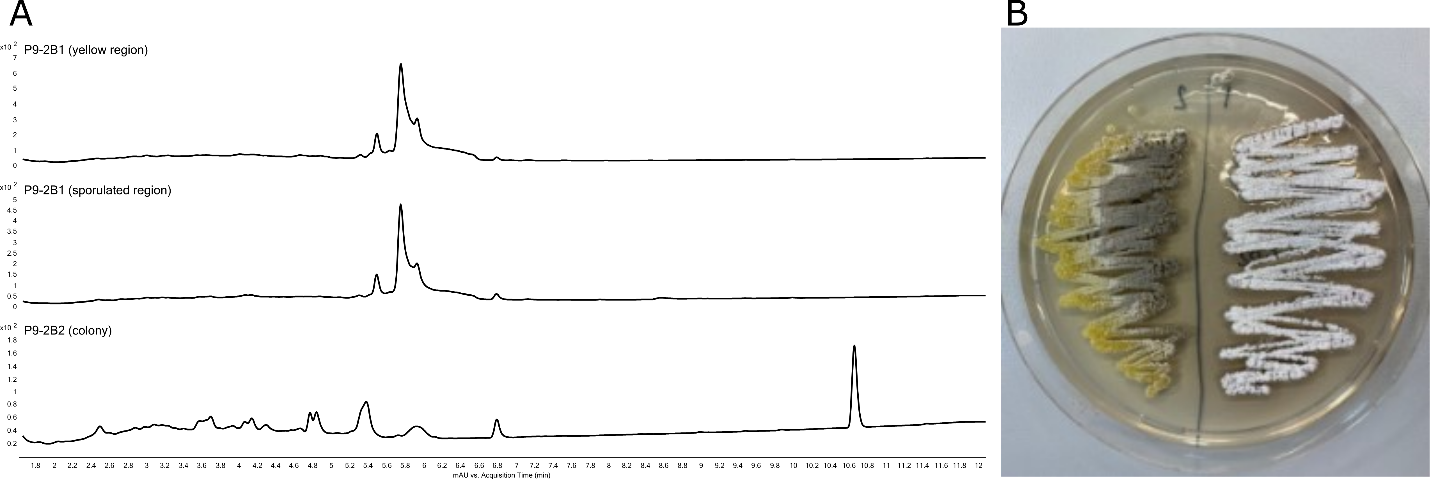

Figure S5. (A) Base peak chromatograms of agar plugs extracted from the regions identified in (B). No discernible difference in metabolome is present in the sporulated region. *Streptomyces* sp. P9-2B2 is on the right side of the agar plate and *Kitasatospora* sp. P9-2B1 is on the left.

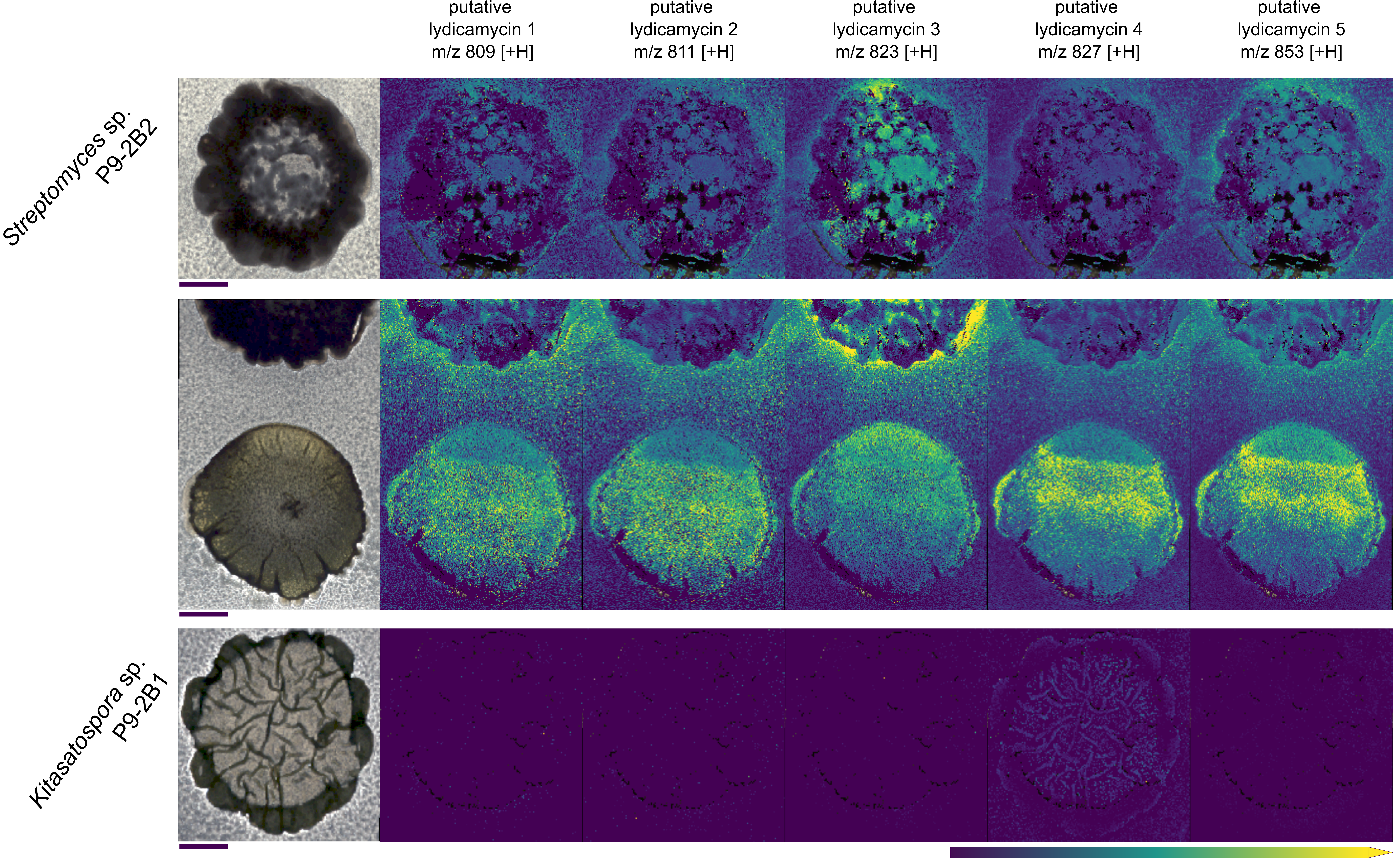

Figure S6. Putative lydicamycins detected via MALDI Mass Spectrometry Imaging. Scale bars: 4mm. Colored scale bar indicates relative ion abundance with hotspot removal activated.

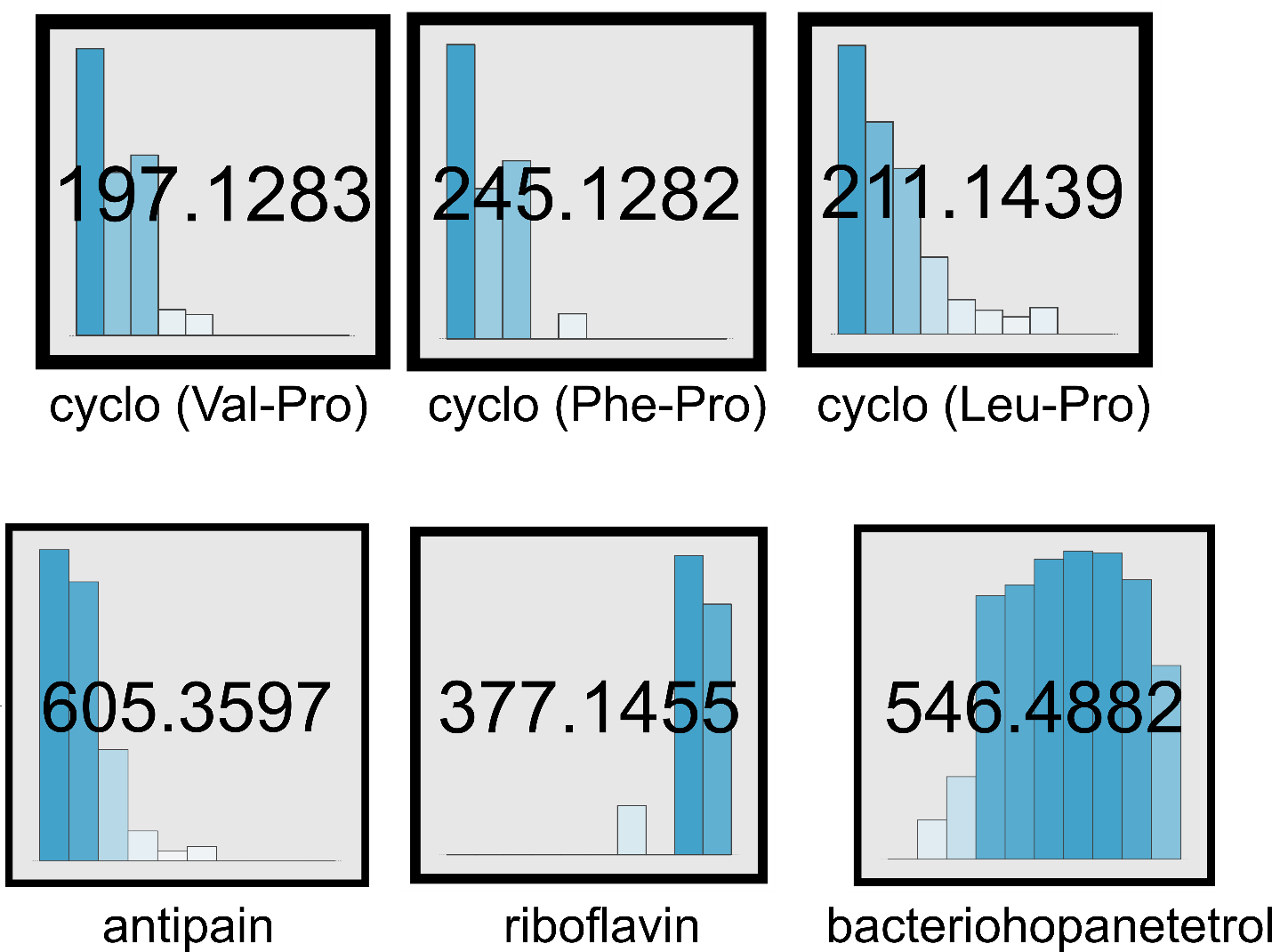

Figure S7. Additional known metabolites identified via feature based molecular networking. Bar graphs within each node is representative of the 10-day time scale which *Streptomyces* sp. P9-2B2 was grown on PDA plates. Coloration of the bar graphs are representative of the peak area (darker = more area).

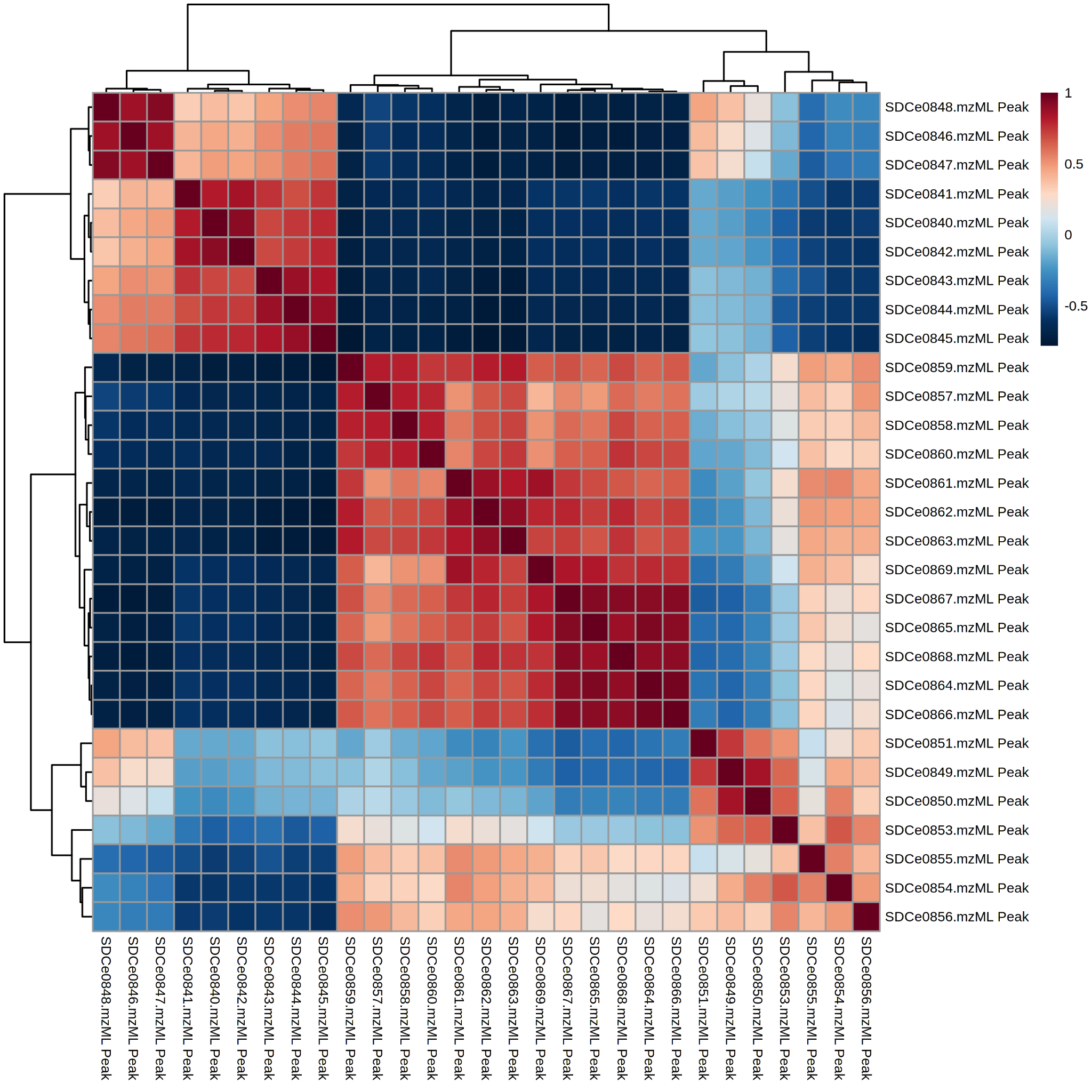
Figure S8. Pearson r correlation of temporal metabolomics samples over 10 days grown on PDA. Sample IDs range from SDCe0840 (Day 1) – SDCe0869 (Day 10), with biological triplicates represented every three sequential samples (*i.e.* 0840-0842 correspond to Day 1 extracts).

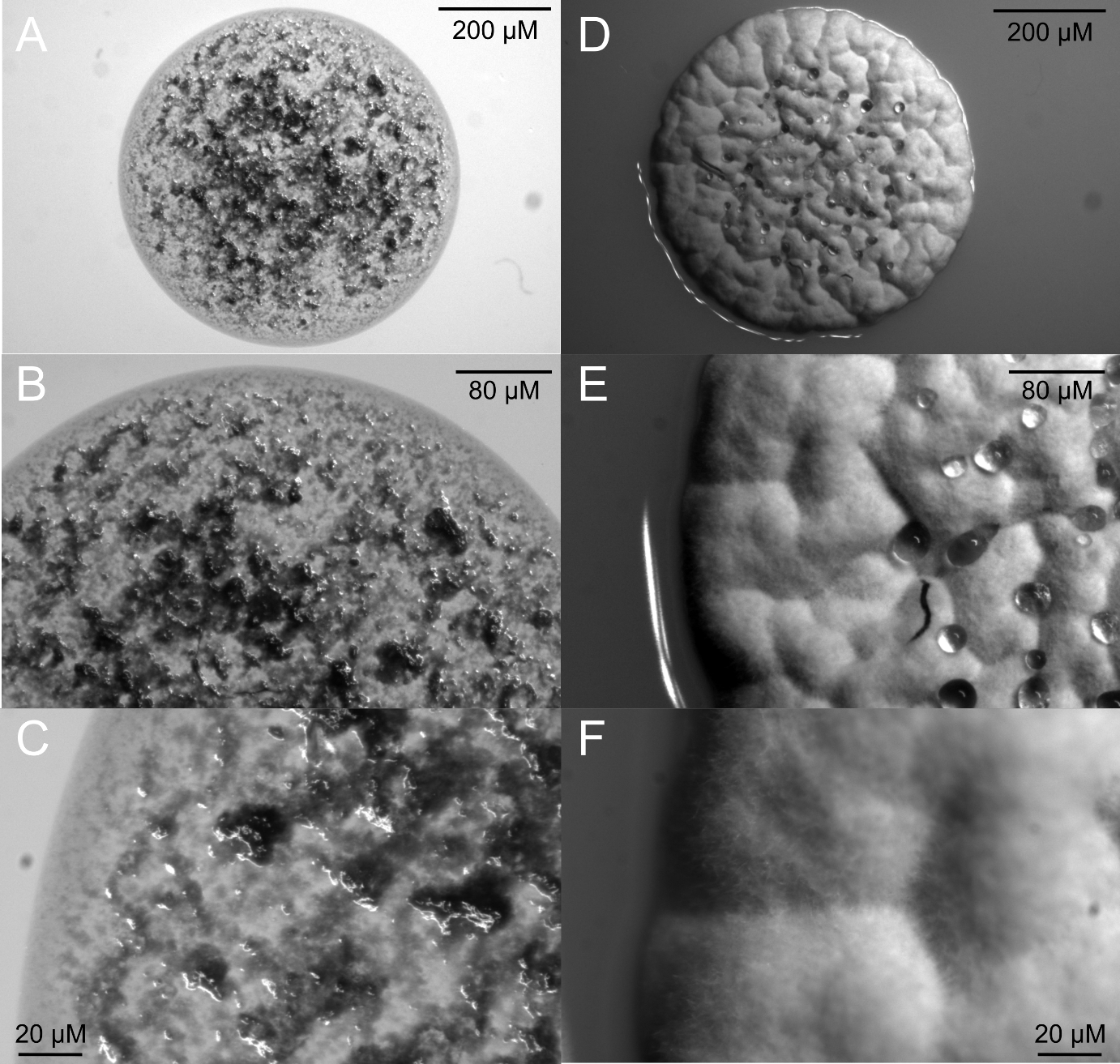

Figure S9. Blank microscopy images of *Streptomyces* sp. P9-2B2 grown on ISP2 (A – 11x, B – 24x, and C – 63x) and PDA (D – 11x, E – 24x, and F – 63x) agar plates for 2 days. On ISP2, pigmentation is clearly visible as well as a ‘wrinkly’ phenotype while on PDA, aerial mycelium formation has begun, with filaments visible (F) and droplets (D and E) due to the hydrophobic layer.

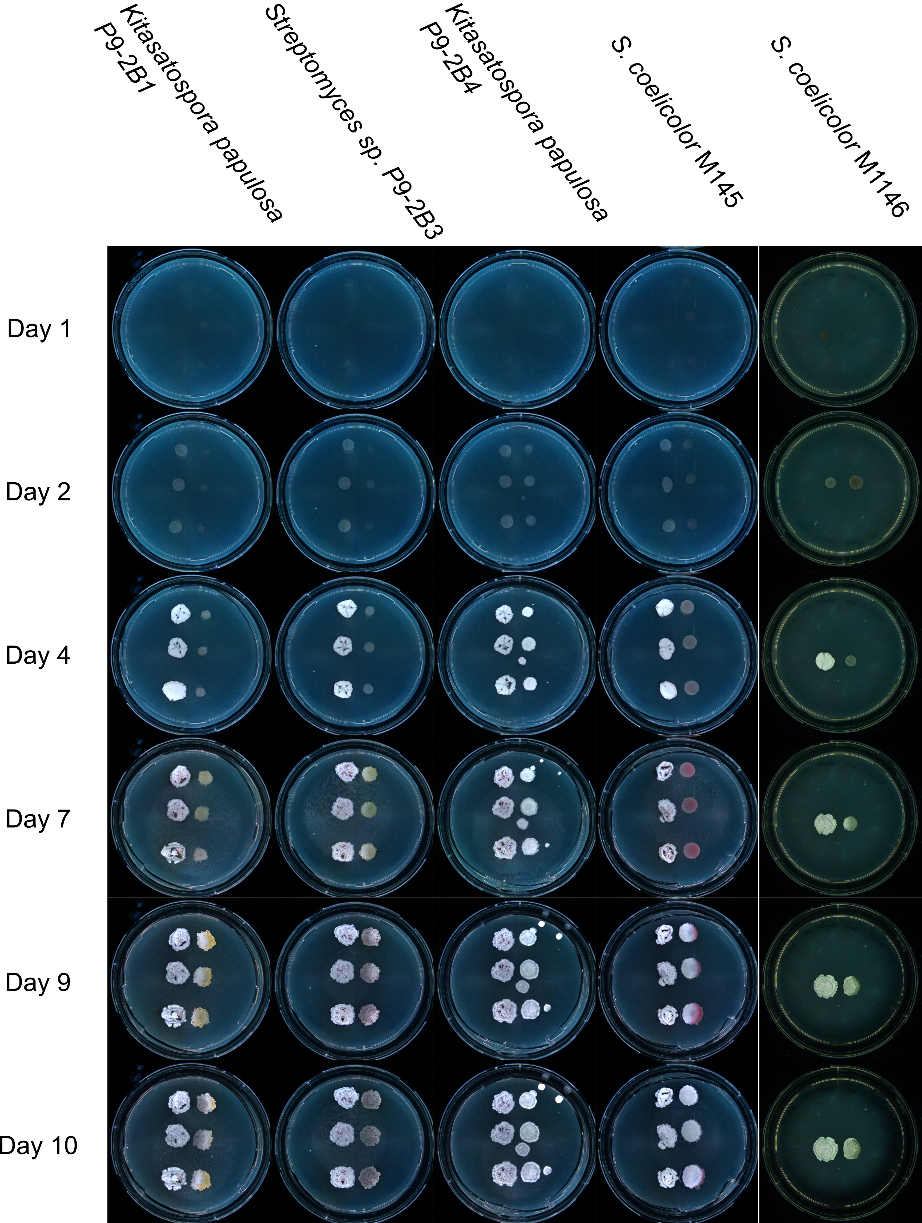

Figure S10. Timelapse microscopy images taken every 24 hours showing the cocultivation of *Streptomyces* sp. P9-2B2 alongside additional environmental isolates and model strain *Streptomyces coelicolor* M1446.

**A**
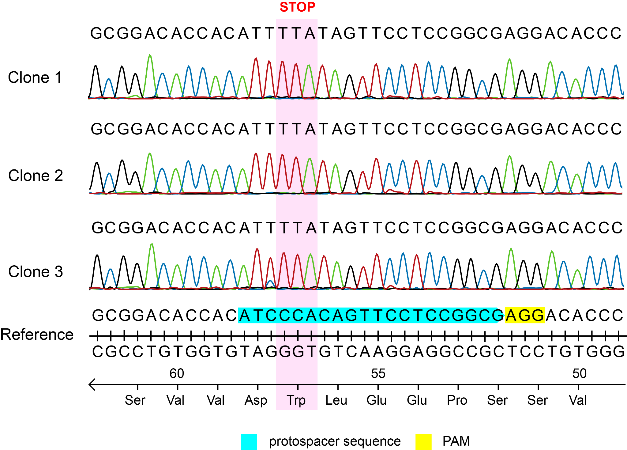
**B**
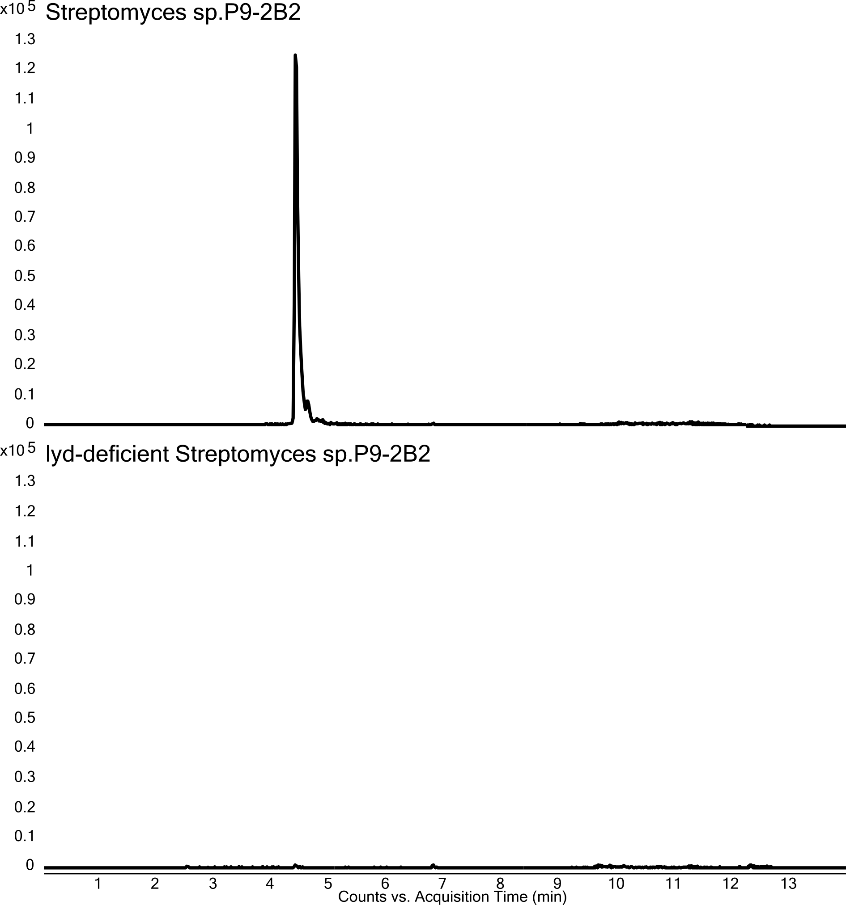

Figure S11. (A) Sanger sequencing of CRISPR base editing application of STOP codon introduction targeting the *lyd60* of *Streptomyces* sp. P9-2B2 strain. The 20-nt protospacer sequence (light green) and the 3-nt PAM sequence (yellow) are shown. (B) Lydicamycin (*m/*z 855.5)) Extract Ion Chromatograms of WT P9-2B2 (top) and *lyd*-deficient P9-2B2 (bottom).

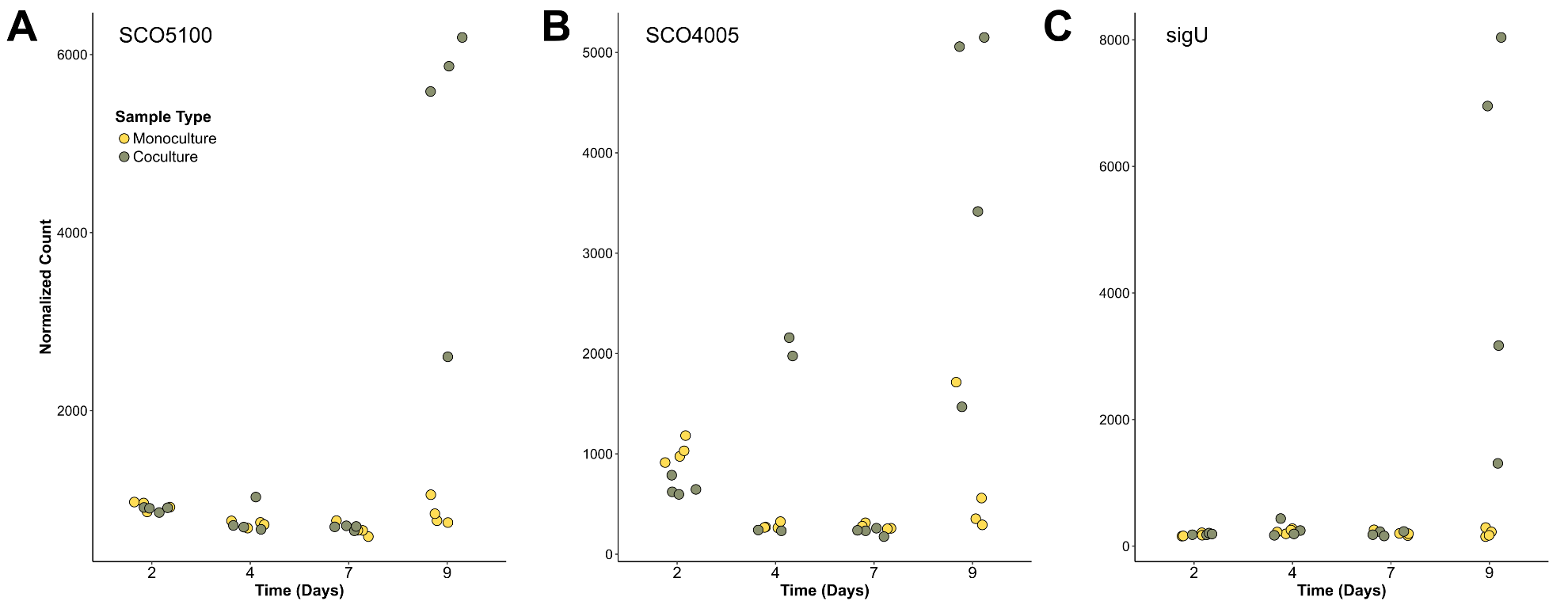

Figure S12. Differential expression of genes tightly associated with cell envelope stress (A) SCO5100, (B) SCO4005 and (C) sigU in *Streptomyces coelicolor* M1146. Yellow data points represent monoculture samples and green represents coculture samples.
